# Viral diversiy within marine biofilms and their potential roles in microbially influenced corrosion

**DOI:** 10.1101/2024.06.11.598143

**Authors:** Chengpeng Li, Yimeng Zhang, Shuqing Jiang, Wenqing Shi, Yongyi Peng, Yingchun Han, Xiyang Dong, Ruiyong Zhang

## Abstract

In marine environments, a wide variety of microbes exert an important influence on the corrosion of materials. Viruses are widely distributed in biofilms formed on materials and may influence the corrosion process through interactions with key corrosive microbes. However, our understanding of the viral communities within biofilms and their implications for corrosion remains notably limited. To fill this knowledge gap, we utilized 53 publicly metagenomes to investigate the ecological dynamics of viruses within biofilms on 8 different materials and their potential roles in the corrosion of materials. In biofilms, the type of materials may be the primary factor driving differences in the diversity of viral communities. Viruses within biofilms predominantly belong to *Caudoviricetes,* and phylogenetic analysis of *Caudoviricetes* and protein-sharing networks with other environments revealed the presence of numerous novel viral clades in biofilms. The presence of virus‒host linkages revealed a close association between viruses and corrosive microbes in biofilms. Viruses play a key role in corrosion by modulating host corrosion-related metabolism. It was observed that viruses could enhance host resistance to metals and antibiotics via horizontal gene transfer. Furthermore, viruses can protect themselves from host antiviral systems through anti-defense systems. Overall, this work illustrates the diversity of viruses within biofilms formed on materials and the intricate interactions between viruses and corrosive microbes, revealing the potential roles of viruses in microbially influenced corrosion.

## Introduction

Microbially influenced corrosion (MIC) refers to the corrosion of materials such as metals and concrete due to the activity of microbes [1]. The MIC process is widespread across numerous industries, including oil and gas, ship, marine structure, and medical industries, and significantly impacts both the economy and safety [2–5]. In China alone, the cost of corrosion in 2014 was estimated to be approximately 310 billion USD, representing 3.34% of the gross domestic product (GDP) for that year [6]. Biofilms, aggregations of microbes and extracellular polymeric substances predominantly composed of polysaccharides, proteins, and extracellular DNA, serve as the main site of MIC [7]. Corrosive microbes can adhere firmly to material surfaces through biofilms and interact with materials [8]. Biofilms create a stable environment for microbes, aiding them in resisting external antimicrobial agents and environmental pressures [9].

A diverse and intricate array of MIC mechanisms exist within corrosive biofilms. Sulfate-reducing, nitrate-reducing, and iron-oxidizing prokaryotes are well known to cause corrosion. Sulfate-reducing *Desulfovibrio vugaris* can accelerate the corrosion of carbon steel through extracellular electron transfer (EET) and metabolite corrosion. Specifically, EET involves transferring electrons generated from the oxidation of extracellular iron across the cell wall into the cytoplasm, where they are used to reduce the external electron acceptor sulfate [10]. Similarly, nitrate-reducing *Pseudomonas aeruginosa* can couple the reduction of nitrate with the oxidation of extracellular iron, accelerating the corrosion of carbon steel [11]. Additionally, iron-oxidizing *Acidithiobacillus ferrooxidans* can accelerate the corrosion of carbon steel by facilitating the oxidation of Fe^2+^ [12]. The metabolic activities occurring within biofilms can alter local microenvironments, such as dissolved oxygen levels, pH fluctuations, organic acid production, and other parameters, thereby significantly influencing electrochemical potentials and indirectly affecting the state and kinetics of corrosion [8]. However, the relationship between viruses and MIC mechanisms remains unclear.

Viruses are among the most abundant and diverse biological entities on Earth [13]. In ecosystems, viruses play important roles. (i) Viruses can impact the community size of prokaryotes through lysis [14]. (ii) Viruses promote the metabolic functions of their hosts through the provision of auxiliary metabolic genes (AMGs) [15]. (iii) Viruses mediate horizontal gene transfer (HGT), altering the host genome by transferring genetic material from the virus to the host. This may result in the host acquiring new traits, such as antibiotic resistance or metal resistance [16, 17]. Additionally, in the process of virus‒host interactions, prokaryotes have evolved antiviral systems such as CRISPR‒Cas systems, restriction-modification (RM) systems, and abortive infection (Abi) systems to defend against viral predation [18]. Moreover, viruses have also evolved anti-defense mechanisms, such as anti-restriction, anti-CRISPR, and other proteins, facilitating coevolution with their hosts [19, 20].

Although the ecological functions of viruses in global ecosystems have been widely explored, the role of viruses in MIC has yet to be clearly recognized. Research on MIC in natural environments has primarily focused on exploring the microbial composition of corrosive biofilms and their potential contributions to corrosion [21]. In marine environments, there is a rich and vast diversity of viruses in biofilms that engage in complex interactions with prokaryotes [22, 23]. These interactions can significantly impact the stability of biofilms and the dynamics of prokaryotic communities and may further affect MIC. In this study, we conducted a comprehensive analysis of viral diversity and phylogeny in corrosive biofilms, explored the impact of viruses on corrosive microbes, and inferred the contributions of viruses to MIC based on these impacts.

## Results

### Overview of prokaryotic communities

To evaluate the prokaryotic community composition across different materials, *rplB* was selected as the marker gene for subsequent analysis. The prokaryotes were predominantly bacteria (92%), with the most abundant phyla being *Pseudomonadota* (39.3% on average), *Desulfobacterota* (9.8% on average), *Planctomycetota* (7.8% on average), and *Bacteroidota* (4.7% on average) (Additional file 2: Table S2 and Additional file 1: Figure S3). The most abundant archaeal phylum was *Thermoproteota* (4.7% on average). The Cs21 sample, which was uniquely derived from laboratory culture, was primarily composed of *Methanobacteriota* (82.4%). *Desulfobacterota* were widespread across the Cs samples (24% on average). Alpha diversity metrics, including Shannon diversity, Simpson diversity, and Chao1 richness, exhibited significant differences among the materials (p < 0.001) (Additional file 1: Figure S4a-c). Principal coordinate analysis (PCoA) was employed to illustrate the distinct community structures associated with different materials (R = 0.36, p = 0.001) (Figure S4d).

The assembly and binning of metagenomes, followed by clustering at 95% average nucleotide identity (ANI), led to the generation of 1006 medium- or high-quality prokaryotic metagenome-assembled genomes (MAGs). The 963 bacterial MAGs and 43 archaeal MAGs recovered from 53 metagenomic samples spanned 65 phyla (Figure S5). The domain Bacteria was primarily composed of *Pseudomonadota* (n = 336), *Planctomycetota* (n = 109), and *Bacteroidota* (n = 69) (Table S3 and Figure S5). The dominant lineages within the domain Archaea were *Halobacteriota* (n = 11) and *Thermoproteota* (n = 19). Notably, only the phylum *Pseudomonadota* was detected in all the materials (Table S3). Some phyla exhibited material sample specificity; for instance, *Dependentiae* was identified only in Rc, and *Thermoplasmatota* was identified solely in Cs.

### Viral communities in biofilms

A total of 18,721 potential viral contigs (>10 kb) were identified, retaining 1,801 viral contigs with completeness greater than 50% (Table S4, Fig. 2a). The remaining contigs were then clustered into species-level viral operational taxonomic units (vOTUs) at 95% ANI, resulting in a total of 1,376 species-level vOTUs.

For taxonomic classification, vOTUs were assigned to taxonomic ranks by searching for marker genes, comparing viral sequences from the Integrated Microbial Genome/Virus (IMG/VR) database, clustering with NCBI RefSeq and constructing phylogenetic trees of *Caudoviricetes*. The vast majority of vOTUs could be annotated at the phylum level, predominantly including *Uroviricota* (n = 1354) (Fig. 1b). *Caudoviricetes* was the only class annotated within *Uroviric*ota. In *Caudoviricetes*, 2.7% of vOTUs were annotated to 11 families at the family level, including *Aut*ographiviridae (n = 14), *Winoviridae* (n = 4), *Peduoviridae* (n = 3) and others (Fig. 1b). Regarding viral lifestyles, 444 vOTUs were identified as lytic, and 210 vOTUs were identified as lysogenic (Fig. 2b and Table S5). There were 34 archaeal viruses and 392 phages (Fig. 1b).

**Fig.1.**
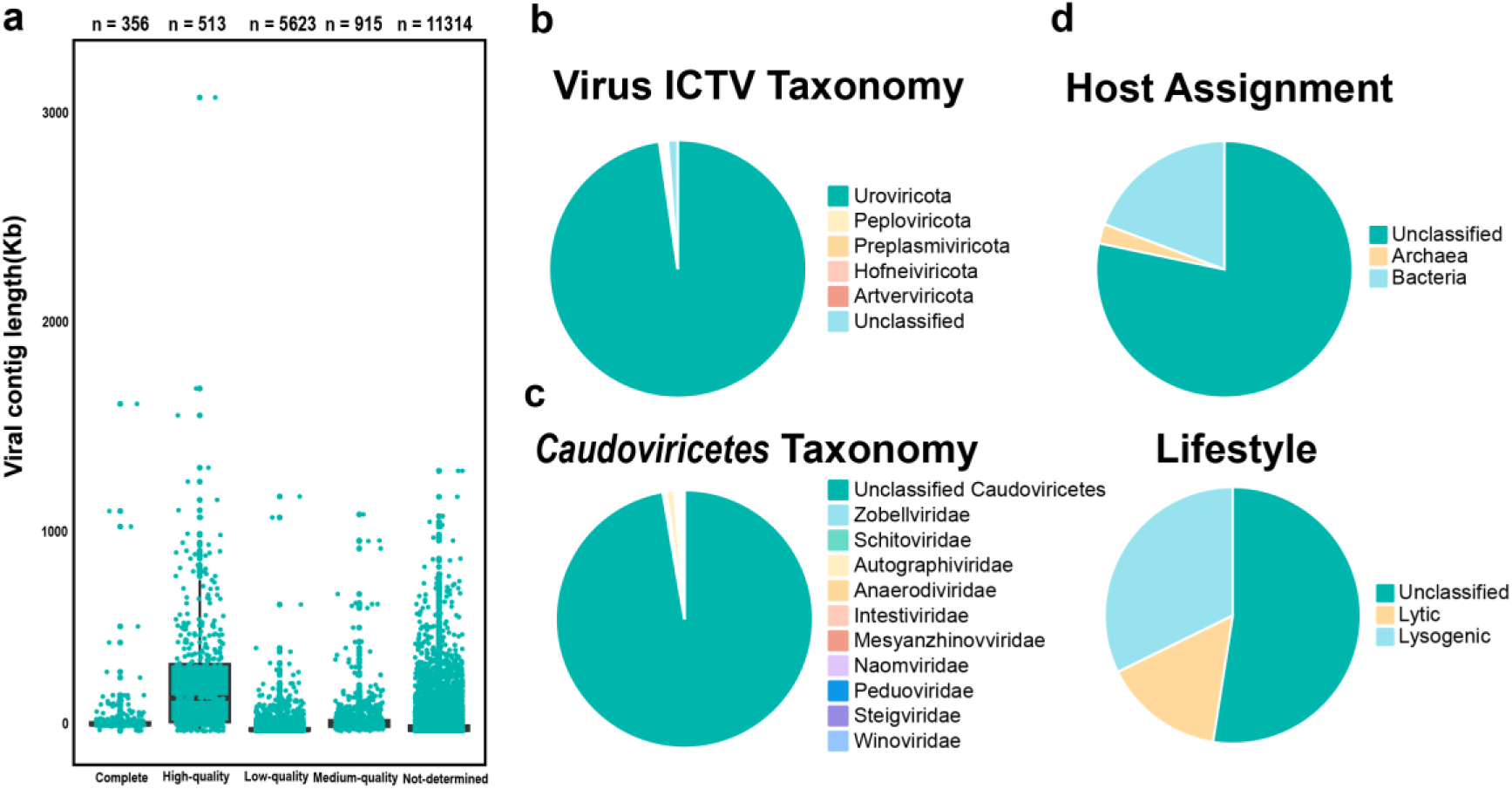
Features of viruses from different materials. **a** Boxplot of contig lengths and quality of predicted viral contigs assessed by CheckV. **b** Classification of viruses according to International Committee of Viral Taxonomy (ICTV) at phylum level. c Classification of *Caudoviricetes* according to ICTV at family level. **d** Host assignment and lifestyle of viruses.

**Fig.2.**
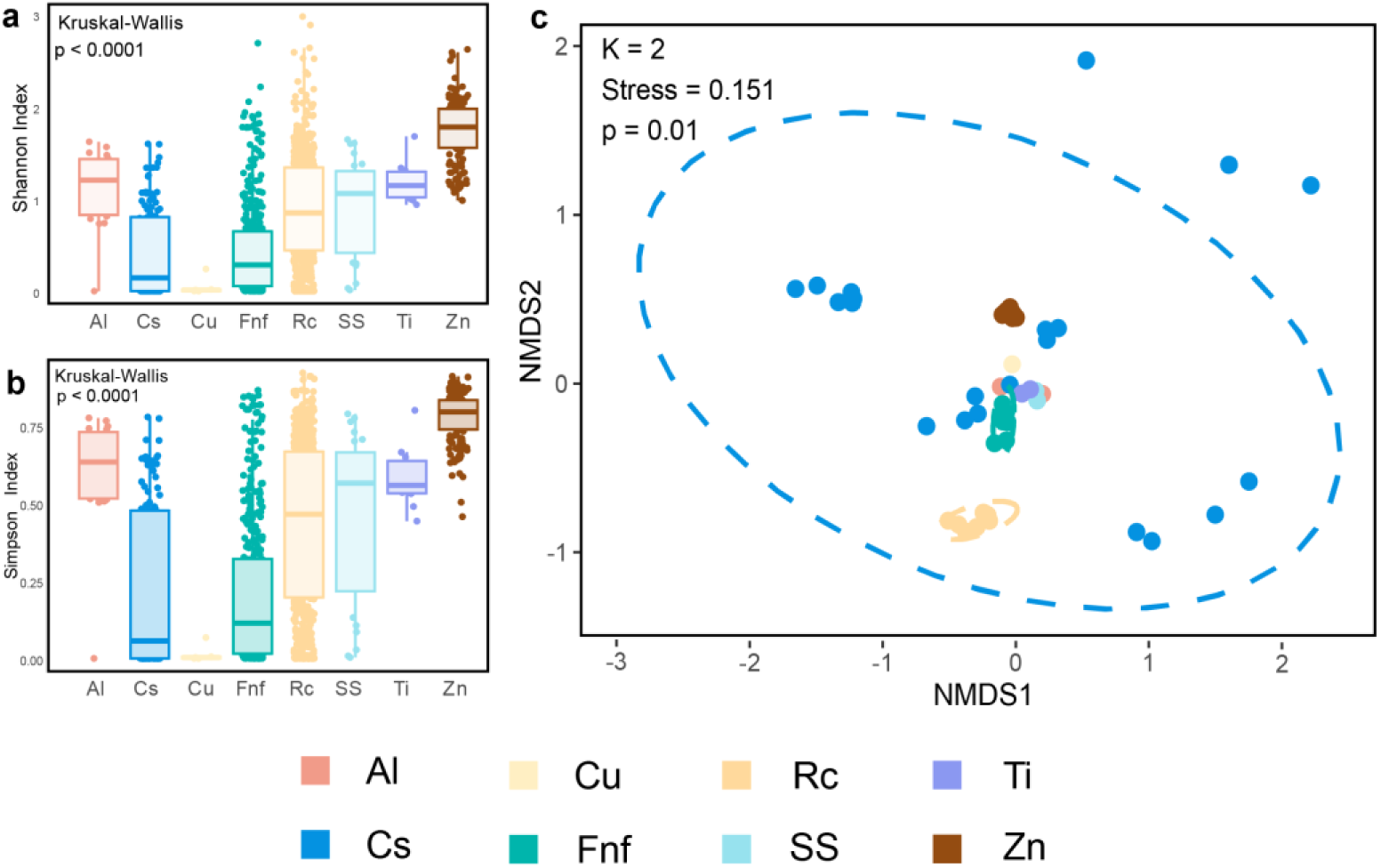
Comparison of viral community diversity between different materials. **a** Shannon indices of viral community diversity from different materials. **b** Simpson indices of viral community diversity from different materials. **c** NMDS analysis of a Bray-Curtis dissimilarity matrix calculated from RPKM values of vOTUs. ANOSIM was applied to test the difference in communities between different materials.

The analysis of alpha diversity revealed significant differences in Shannon and Simpson diversity among the various materials (p < 0.0001) (Table S6). Viruses from the Zn treatment group exhibited the greatest Shannon and Simpson diversity of viral communities (Shannon: 1.7 ± 0.3, Simpson: 0.8 ± 0.1). Nonmetric multidimensional scaling (NMDS) analysis revealed that there were significant differences in the viral communities among different materials (R = 0.35, p = 0.001) (Fig. 2).

*Caudoviricetes*, known as tailed viruses, are widely distributed across diverse ecosystems [24]. To explore the distribution and taxonomy of *Caudoviricetes* within biofilms, a phylogenetic tree comprising 97 vOTUs was constructed using 77 marker genes. The data revealed that *Caudoviricetes* were distributed across five materials, with the majority of viruses not being classified at the family level (Fig. 3a). To achieve a more effective taxonomic classification of *Caudoviricetes* members, NCBI RefSeq and viral sequences from this study were used to construct a phylogenetic tree based on the 77 marker genes of *Caudoviricetes* (Fig. 3b). However, a significant majority of *Caudoviricetes* (92.8%) remained unclassified, indicating a substantial presence of unknown viral clades within biofilms.

**Fig.3.**
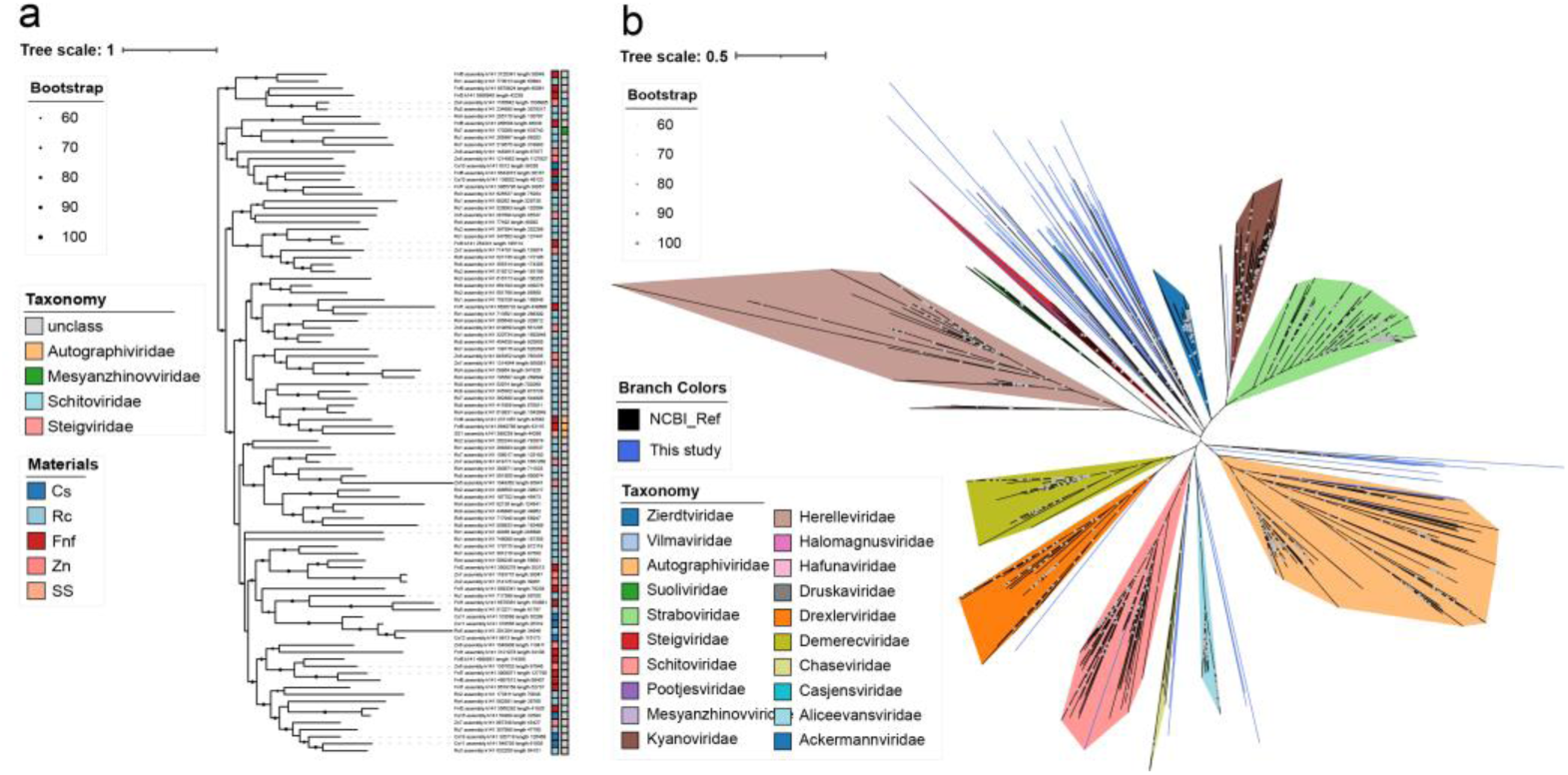
Phylogenomic tree of the vOTUs from the class *Caudoviricetes.* **a** A maximum-likelihood phylogenetic tree of 97 species-level vOTUs from different materials. **b** A maximum-likelihood phylogenetic tree of 1071 species-level vOTUs from different materials and NCBI reference.

To further explore viral communities in biofilms formed on materials and their relationships with viral sequences from other environments, a gene-sharing network was constructed using vConTACT2. In this network, each viral cluster was considered equivalent to a genus-level taxonomic classification. Viral clusters from biofilms, acid mine drainage, seawater, soil, and cold seeps formed 27,218 viral clusters (VCs) (Table S7 and Figure S6b). No VCs were found to exist across all ecosystems. Viruses from the GOV2.0 database formed the greatest number of unique VCs (n = 21,890). The 1,376 vOTUs from biofilms were clustered into 548 VCs, with 62.6% of these VCs shared with VCs from other environments. Among these 548 VCs, a significant number (n = 297) of viral clusters from biofilms shared VCs with viruses according to the GOV2.0 database, indicating high similarity between viral communities in biofilms and those in marine environments. Additionally, 57 VCs of biofilm viruses were shared with cold seeps, 39 with acid mine drainage, 34 with soil, and 28 with NCBI RefSeq. In addition to VCs shared with other environments, the presence of 37.4% of VCs exclusively in biofilms reveals the endemism of biofilm viruses.

### Virus-host linkages

To investigate the potential influence of viruses on prokaryotic communities across various materials, a variety of methods were employed to predict in situ and ex situ virus‒host linkages (Table S8 and Fig. 4a). In total, 421 potential in situ virus‒host linkages were identified, of which 294 vOTUs were linked to 231 prokaryotic MAGs (Fig. 4b). These hosts were distributed across 26 bacterial phyla and 3 archaeal phyla. Most of the predicted host MAGs were assigned to 3 bacterial phyla, *Pseudomonadota* (n = 110), *Planctomycetota* (n = 99), and *Nitrospinota* (n = 36), while the most frequently infected archaeal phylum was *Thermoproteota* (n = 24). Of the virus‒host linkages identified, 214 vOTUs (72.7%) were predicted to have a narrow host range, whereas 80 vOTUs (27.3%) were predicted to infect multiple hosts. The analysis of viral and host communities across various materials revealed a wide range of host phyla present in Cs, Fnf, Rc, and Zn (Figure S7). In Cs and Rc, the proportion of viruses with lysogenic cycles exceeded 50%.

**Fig.4.**
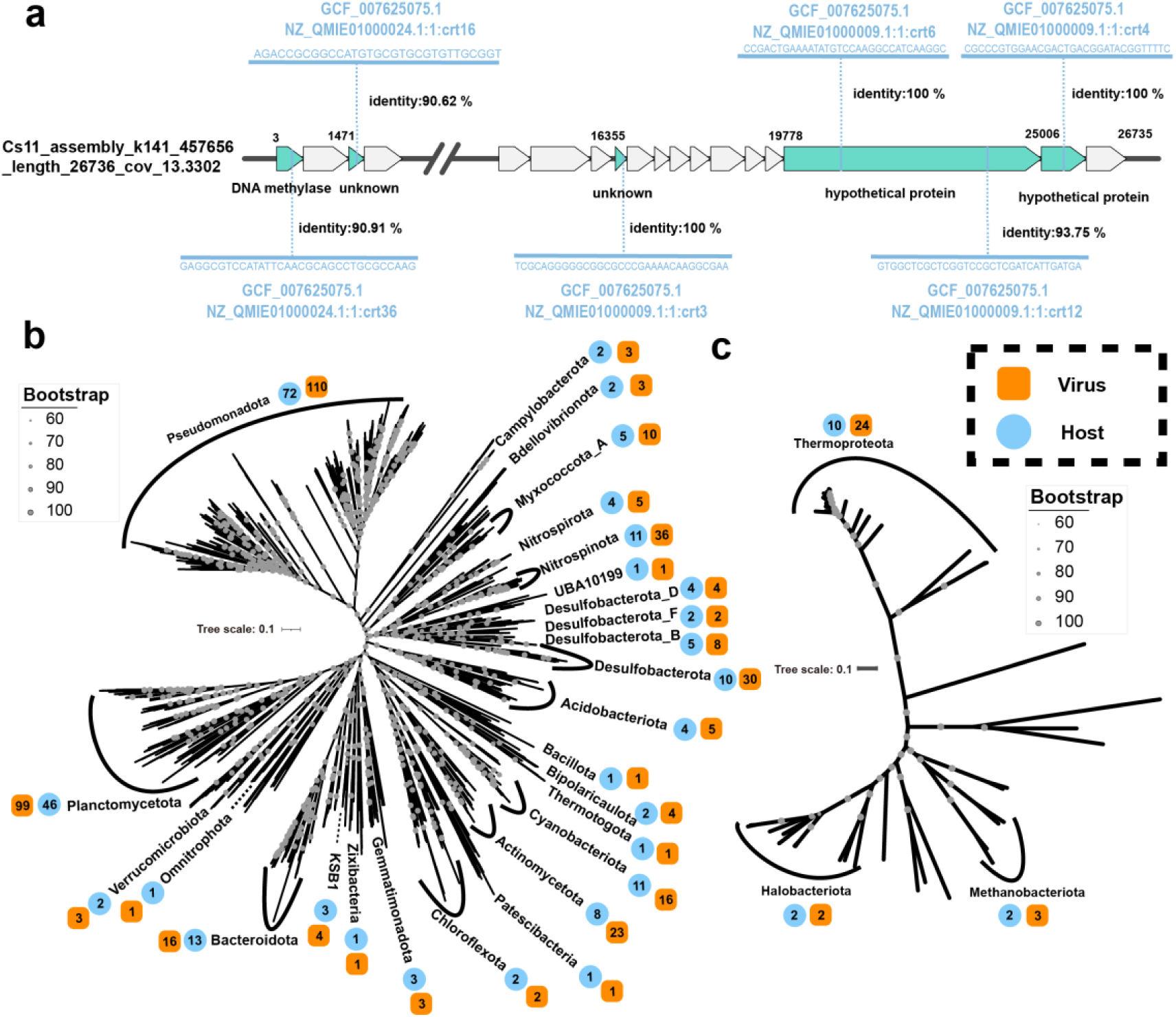
Virus-host linkages in the biofilms. **a** Virus-host linkages based on CRISPR spacers. CRISPR spacers in viral genome and host (GCF_007625075.1) are shown in blue. Maximum-likelihood phylogenetic trees of bacterial (**b**) and archaeal (**c**) MAGs at the phylum level were inferred from a concatenated alignment of bacterial or archaeal single-copy marker genes. The number of hosts in each phylum is shown in blue, while the number of viruses infecting hosts is shown in orange.

To investigate the potential impact of viruses on the ecology of prokaryotic communities within biofilms, the infection dynamics of viruses to their hosts were assessed based on the relative abundance of viruses and hosts. Among viruses with a lysogenic cycle, there was a positive correlation (r = 0.53, p < 0.001) between viral abundance and host abundance (Figure S8). A negative relationship was observed between viral abundance and the virus‒host abundance ratio, suggesting that viruses with lysogenic cycles were more abundant in dominant hosts, which supports the Piggyback-the-Winner (PtW) hypothesis [25]. This indicates that the phenomenon of superinfection exclusion occurs within biofilms on materials, where hosts with high density tend to recruit lysogenic viruses as a defense mechanism against infection by closely related viruses. Among viruses with lytic cycles, the virus infecting the corrosive microbe Cs19_bin.2 was selected as an example to show its genomic context and structural protein. The results indicate that structure-based functional prediction can broaden the scope of viral protein annotation (Figure S8 and Table S9). The docking results revealed a tight interaction (confidence score = 0.9435, RMSD = 43.58) between the major capsid protein and portal protein of viruses infecting Cs19_bin.2 (Table S10).

### Potential impact of AMGs, ARGs and MRGs on corrosion-related microbes

In this study, we focus on the interactions between viruses and three commonly studied corrosion-related microbes (Fig. 5 and Table S11): (i) Sulfate-reducing prokaryotes (SRP): SRP can accelerate corrosion by utilizing electrons generated from the oxidation of Fe^0^ to Fe^2+^ for dissimilatory sulfate reduction. This is currently the most extensively studied corrosive microbe [26]. (ii) Nitrate-reducing bacteria (NRB): NRB can accelerate corrosion by utilizing electrons generated from the oxidation of Fe^0^ to Fe^2+^ for dissimilatory nitrate reduction. This corrosion mechanism is similar to that of SRP [11]. (iii) Iron-oxidizing bacteria (IOB): These bacteria not only accelerate corrosion through the oxidation of Fe^2+^ to iron but also induce environmental acidification. This acidification process significantly accelerates the corrosion and deterioration of concrete structures [27, 28].

**Fig.5.**
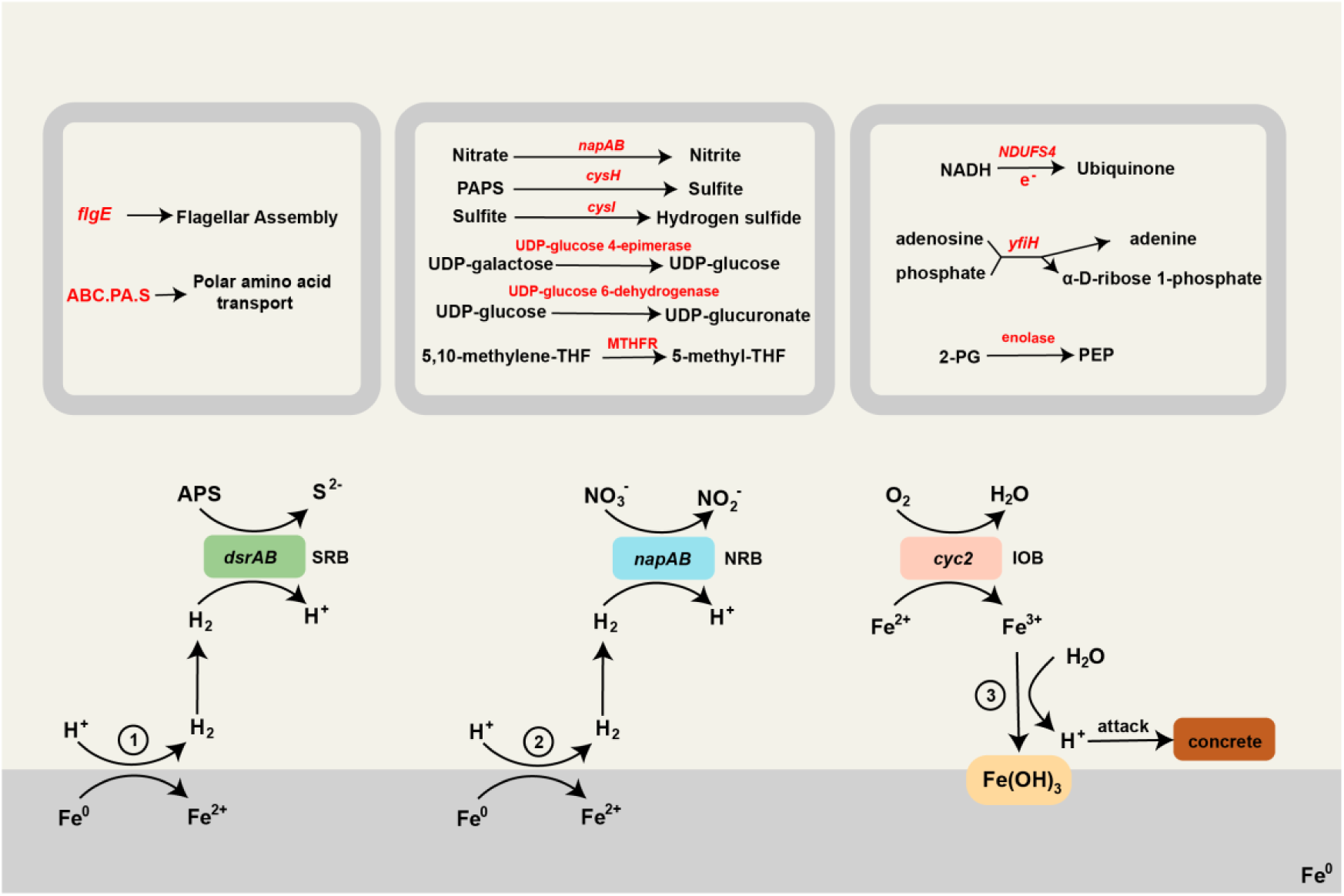
AMGs of viruses infected on corrosive microbes. AMGs-involved pathways is shown in box. AMGs is highlighted in red. Figures show three MIC mechanisms of potential corrosive microbes SRB, NRB, and IOB. The corrosion-related genes in SRB, NRB and IOB are shown in green, blue and pink.

**Fig.6.**
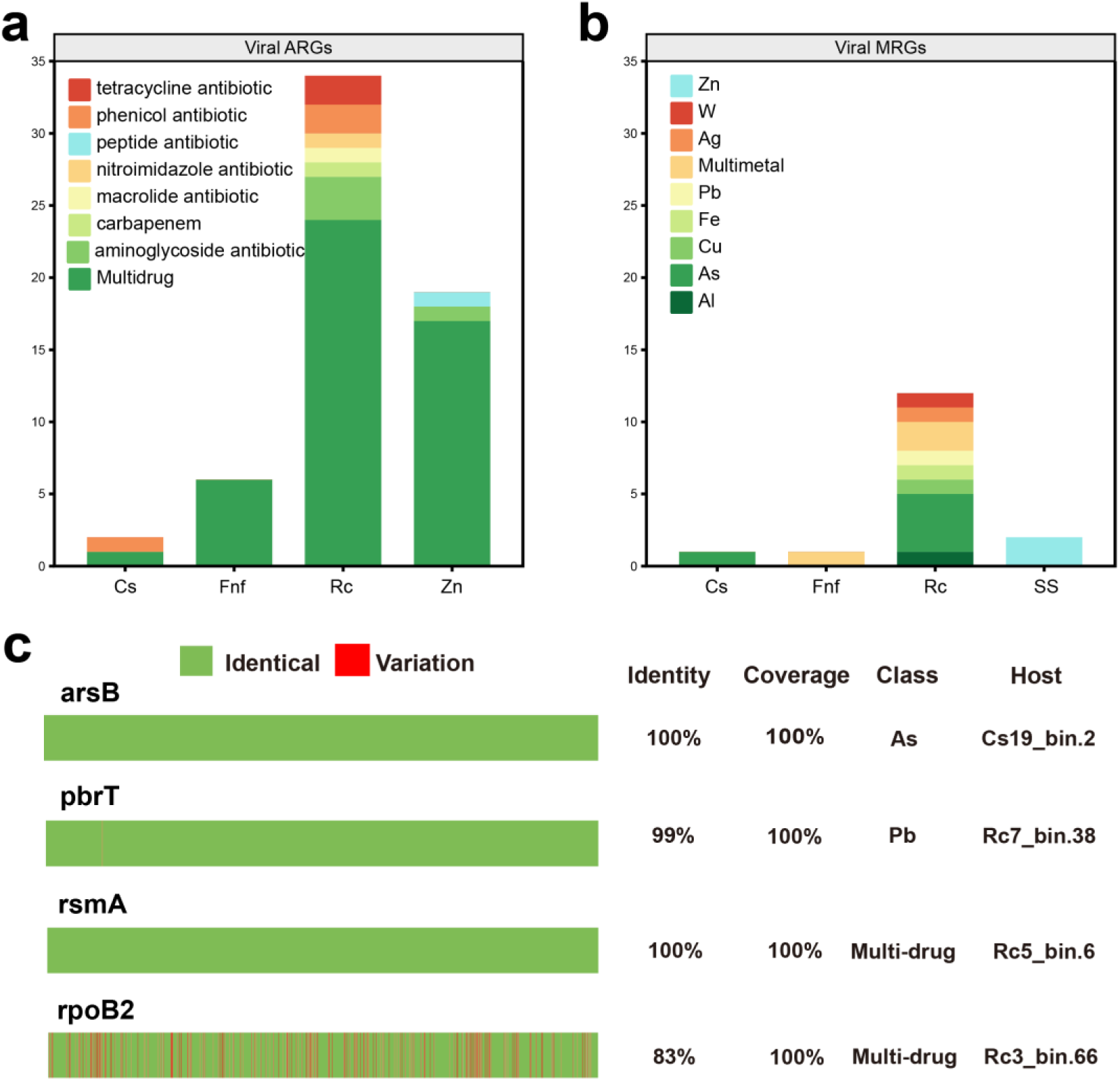
Resistant drugs, metals and sequence alignments of viral ARGs and MRGs. **a** Number of viral ARGs among different materials. **b** Number of viral MRGs among different materials. **c** Comparison of viral ARGs and MRGs with host genomes, variable sites are in red. The coverage and identity are based on blastn algorithm.

To investigate the potential role of viruses in biofilms, a total of 77 AMGs were selected for subsequent analysis (Table S12). Based on the functional categories annotated by DRAM-v, these AMGs were classified into four main groups: carbon Utilization (n = 12), energy (n = 26), miscellaneous (MISC) (n = 12), and transporters (n = 26). Notably, viruses carrying 43 AMGs were associated with specific prokaryotic hosts.

Among the identified AMGs, *napA* and *napB*, along with *NDUFS4*, were found to have potential associations with the MIC. The AMGs *napA* and *napB* play critical roles in the reduction of nitrate to nitrite. NapA facilitates this process by receiving electrons from NapB, while NapB, a key electron transfer subunit in the periplasmic nitrate reductase complex, can accept electrons from NapC. Viruses harboring AMGs encoding *napA* and *napB* were predicted to infect the NRB Rc7_bin.54 and IOB Rc3_bin.47. This finding implies that viruses may enhance nitrate reduction via the AMGs *napA* and *napB*, thereby increasing the MIC. The phylogenetic tree showed that the levels of NapA from viruses were closely related to those from *Acidobacteria* and *Calditrichota* (Figure S9a). Pairwise structural alignment revealed that the NapA protein from the virus exhibited high similarity to that from *Calditrichota* (TM-score = 0.8378), suggesting potential functional similarities (Figure S10b). The docking results of NapA and NapB revealed a high score (confidence score = 0.9505, RMSD score = 95.37) for the NapAB complex, indicating its potential to assemble into a complex (Table S8). Furthermore, *NDUFS4*, identified as an accessory subunit of the mitochondrial membrane respiratory chain NADH dehydrogenase, plays a role in transferring electrons from NADH to the respiratory chain. This implies that viruses may have the potential to promote the electron transfer of corrosive microbes.

AMGs not only contribute to corrosion-related metabolic pathways in corrosive microbes but also engage in broader metabolic processes (Figure S11). AMG associated with sulfur cycling, including phosphoadenosine 5’-phosphosulfate reductase (*cysH*), was found in viruses infected with the NRB Rc7_bin.54 The *cysH* gene is involved in catalyzing the transformation of phosphoadenosine 5’-phosphosulfate to sulfite, a critical step in sulfur metabolism [29]. AMGs associated with carbon cycling, including UDP-glucose 4-epimerase and UDP-glucose 6-dehydrogenase, was found in viruses infected with the NRB Rc7_bin.54 and IOB Rc3_bin.47. The gene encodes UDP-glucose 4-epimerase, which can facilitate the conversion of UDP-galactose to UDP-glucose, representing the final step in galactose metabolism. Additionally, UDP-glucose 6-dehydrogenase can catalyze the conversion of UDP-glucose to UDP-glucuronate. In addition to metabolic pathways associated with elemental cycling, AMG encodes *flgE*, a component of the bacterial flagellar hook structure essential for motility. The *flgE* gene encodes a part of the flagellar hook, a critical structure that connects the flagellar motor to the filament, playing essential roles in flagellar biosynthesis and biofilm formation [30]. The presence of AMG *flgE* suggested a potential influence on the mobility of SRP, which could affect biofilm formation and consequently MIC processes in marine environments.

In addition to AMGs, viruses are also capable of encoding antibiotic resistance genes (ARGs) and metal resistance genes (MRGs), facilitating their transfer to host genomes. In this study, a comprehensive search against several databases led to the identification of 61 ARGs and 16 MRGs (Fig. 5a-b and Table S13). The predominant types of ARGs included multipledrugs, aminoglycosides, and tetracyclines, whereas the primary MRG types were associated with multimetal and arsenic resistance (Figure S11ab).

To assess the impact of viruses on the propagation of ARGs and MRGs within biofilms, vOTUs encoding these genes were compared with the genomes of their hosts. The homologous genes corresponding to the viral AMGs and ARGs, which exhibit 70% nucleotide identity and no less than 80% coverage, were identified in the host genomes (Fig. 5c and Table S14). A notable example among SRPs is the viral MRG encoding the arsenite resistance gene *arsB* from a Cs sample, which showed high homology to its host genome (Figure S11c). The *arsB gene* confers resistance to arsenite by enabling cells to extrude this compound. The phylogenetic tree showed that *arsB* from the virus was related to *arsB* from *Pseudodesulfovivrio* (Figure S12). Remarkably, the ARG *rsmA,* encoding a regulatory factor of the resistance-nodulation-cell division antibiotic efflux pump, showed high amino acid sequence identity with its host Rc5_bin.6. In summary, the high homology between viral ARGs, MRGs and the host genome suggests that viruses play a significant role in the transfer and spread of these resistance genes among prokaryotic communities.

### Diverse antiviral strategies and anti-defense systems between viruses and corrosive microbes

Prokaryotes within biofilms have evolved antiviral systems to combat viral infections, with a predominance of restriction-modification and CRISPR‒Cas systems observed among prokaryotes (Figure S13a) (Table S15). The RM and cyclic oligonucleotide-based antiphage signaling system (CBASS) were used for all the materials (Figure S13b). Multiple antiviral systems have been identified in corrosive microbes infected by viruses. For instance, the presence of RM Type II, Cas, AbiE, and CBASS II systems was detected in SRP Cs18_bin.13 (Fig. 7b). RM Type II cleaves DNA at specific recognition sites, thereby protecting the host from viral integration [31]; the Cas system provides adaptive immunity by targeting and cleaving foreign genetic elements [32]. To counter antiviral systems, viruses in biofilms have developed anti-defense systems (Figure S13c) (Table S16). For example, within viruses infecting the IOB Rc3_bin.66, the presence of AcrIE4 was detected (Fig. 7a). The I-E type anti-CRISPR, AcrIE4, plays a crucial role during the infection process by protecting phages from suppression mediated by the Cas system [33]. Such interactions imply the intricate battle between viruses and prokaryotes within the microenvironments of corrosive biofilms.

**Fig.7.**
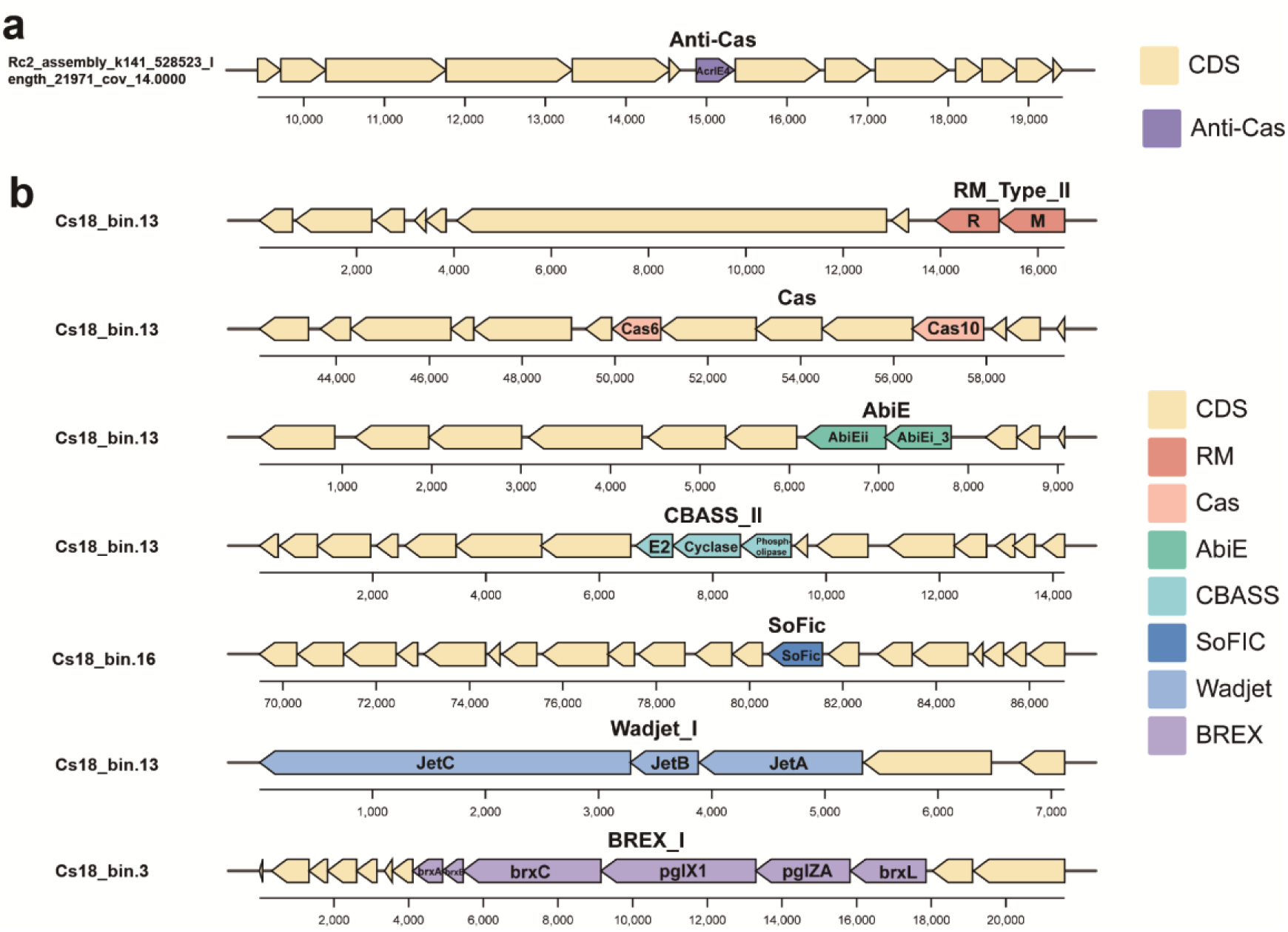
The genomic context of anti-viral system related genes and anti-defense related genes. **a** The genomic context of anti-defense system, CDS region are in yellow. **b** The genomic context of antiviral systems, CDS region are in yellow, anti-viral systems are in different colors.

## Discussion

Viruses may play a critical role in biofilms formed on materials, yet their diversity and functional mechanisms remain insufficiently explored. However, existing studies have investigated the interactions between viruses and their hosts within biofilms, expanding our knowledge of viruses in biofilms [23]. Nevertheless, the potential role of viruses in microbe–material interactions is still unclear. Using 53 public metagenomes, we examined the dynamics of viral communities within biofilms on eight different materials and their interactions with corrosive microbes, revealing the potential impact of viruses on MIC.

In this study, vOTUs were identified across various materials. There were significant differences in viral communities among the materials. Thus, it is supposed that the type of material may be a critical factor influencing viral community composition. Such variations could be attributed to differences in the surface characteristics of the materials, such as surface roughness, chemical composition, and potential differences [34]. This finding aligns with observations from studies in Hong Kong built environments, highlighting the importance of specific materials and surface types as key factors influencing viral diversity [35]. In addition, the diversity of prokaryotes on carbon steel is low, which could influence the diversity of viruses infecting these prokaryotes. This explains why viruses derived from carbon steel exhibit lower diversity. In biofilms, viruses belonging to *Caudoviricetes* dominate the viral communities, suggesting that *Caudoviricetes* largely represent the clades and diversity of viruses within the biofilm. However, the majority of these viruses cannot be classified at the family level, a phenomenon consistent with discoveries in other environments [36]. Compared to other environments, the presence of VCs exclusively in biofilms highlights unique viral clades within biofilms formed on materials.

Viruses interact with their hosts through diverse infection modes. On the one hand, beneficial lysogenic viral infections can shield hosts from new viral infections, enhance host ecological niches, and boost host competitiveness [37, 38]. In marine biofilms formed on materials, viruses with lysogenic cycles constitute more than half of the viral population, and the correlation between the abundance of viruses with lysogenic cycles and host abundance supports the PtW theory. According to the PtW theory, as host density increases, host abundance drives the transition of viral lifestyles from lytic cycles to lysogenic cycles, resulting in a decrease in the viral-host ratio [25]. This phenomenon suggests that viruses with lysogenic cycles in biofilms on materials may enhance the competitive advantage of dominant hosts. On the other hand, viruses with lytic cycles can impact bacterial populations through predation, exerting top-down influences by reducing bacterial numbers and bottom-up effects by breaking down cells [39, 40]. We identified a virus with a lytic cycle that infected a corrosive microbe and revealed key proteins involved in this process. Specifically, proteins such as terminase enzymes and portal proteins play crucial roles in the packaging of viral DNA and the assembly of viral particles [41, 42]. These components not only serve as critical factors for virus replication and maturation but also may further influence the survival and function of the host by regulating the release of viral particles. In the field of MIC, phages are considered to potentially alleviate MIC by lysing corrosive biofilms. The discovery of viruses with lytic cycles infected on corrosive microbes demonstrates the feasibility of this idea. Thus, future studies are required to isolate viruses with lytic cycles in corrosive biofilms [43, 44].

AMGs play a crucial role in regulating host metabolism and microbial-driven biogeochemical cycles [45]. Various AMGs were identified, and two types of AMGs may be associated with MIC. The AMGs *napA, napB* and *NDUFS4* may promote the corrosion of materials by promoting nitrate reduction and electron transfer [46]. AMG *flgE* may be associated with the formation of biofilms in the SRP. In activated sludge, AMGs were found to participate in dissimilatory sulfate reduction [47]. This finding implies that viruses have the potential to directly participate in the MIC process by SRP. Furthermore, viruses can encode other AMGs that enhance the metabolism of corrosive microbes. In Rc, AMG *cysH, which is* involved in sulfur metabolism and has been extensively discovered in other environments, was identified [48, 49]. In our study, we identified AMGs potentially associated with the MIC. Further validation is required to ascertain the actual functions of these AMGs.

Viruses are significant because they can act as carriers of ARGs and MRGs, potentially transferring them to bacteria through horizontal gene transfer [50]. Heavy metals can act as selective pressures on antibiotic resistance and metal resistance, and studies have shown that viruses carry a greater abundance of MRGs and ARGs at high concentrations of heavy metals [16, 51, 52]. However, extracellular polymeric substance can act as a physical barrier, hindering the diffusion and ingress of bacteriophages into the biofilm matrix, thereby reducing the transfer of ARGs and MRGs from viruses [53]. In this research, many viral ARGs and MRGs were identified within the biofilms formed on the materials, indicating that viruses can act as potential reservoirs of ARGs and MRGs. We found that the majority of viral ARGs and MRGs originated from Rc. These parameters are most likely correlated with the highest number of vOTUs from Rc. The tracking of viral ARGs and MRGs in response to corrosive microbes implies that viruses contribute to increasing host survival and adaptation capabilities within biofilms. For example, it was observed that viruses can enhance host resistance to arsenic in biofilms formed on carbon steel.

Frequent interactions between viruses and hosts have led to the evolution of antiviral and defense systems [54]. Prokaryotes can resist viral infections through various antiviral mechanisms. We identified abundant antiviral systems such as the RM, Cas, AbiE, and CBASS systems in corrosive microbes, indicating the presence of diverse mechanisms to counter viral infection in corrosive microbes. Corresponding to antiviral systems, viruses infecting corrosive microbes can evolve anti-defense systems such as the anti-CRISPR system. These findings reveal intimate interactions between viruses and corrosive microbes within biofilms formed on materials.

## Conclusion

This study revealed the diversity, taxonomy, and lifestyles of viral communities within biofilms formed on materials. Novel viruses are associated with prokaryotic hosts, including corrosive microbes. We found that viruses carried AMGs associated with MIC, highlighting the potential role of viruses in MIC. In addition, we found that viruses can act as HGT vectors to spread ARGs and MRGs to corrosive microbes, suggesting that viruses can promote the resistance of hosts in biofilms. Finally, we described antiviral systems and anti-defense systems in corrosive microbes and viruses, indicating coevolution between corrosive microbes and viruses. In marine environments, viruses are crucial elements of biofilms, and a deeper exploration of viruses could significantly enhance our understanding of the ecological role of viruses and viral contributions to MIC.

## Materials and methods

### Metagenomic data collection

The datasets of 53 publicly metagenomes were collected from the National Center for Biotechnology Information-Sequence Read Archive (NCBI-SRA) [21, 22, 27, 55–60] and were specifically chosen for biofilms formed on materials. These samples were categorized into eight different types of materials: carbon steel (Cs, n = 21), aluminum (Al, n = 3), zinc (Zn, n = 9), titanium (Ti, n = 2), copper (Cu, n = 1), stainless steel (SS, n = 2), reinforced concrete (Rc, n = 8), and ferromanganese nodule fields (Fnf, n = 7). The Cs21 sample was only derived from the laboratory. Detailed descriptions and categorizations of these samples are available in Table S1 and Figure S1. An overview of the workflow used for viral analysis in this study is available in Figure S2.

### Composition and diversity of prokaryotic community

To explore the composition and diversity of the prokaryotic community, metagenomic reads were processed with singleM v0.13.2 [61] (parameters: default). *rplB* was selected as the marker gene for defining operational taxonomic units (OTUs). Then, the OTUs were summarized by rarefaction and clustering according to the SingleM protocol (https://wwood.github.io/singlem/).

### Metagenome assembly and generation of metagenome-assembled genomes

The raw reads were processed for quality control with fastp v0.23.2 [62] (parameters: default). Clean reads were individually assembled using MEGAHIT v1.1.3 (k list: 21, 29, 39, 59, 79, 99, 119) [63] within the metaWRAP v1.3.2 pipeline.

For the processing of each assembly, contigs (> 1000 bp) were subjected to a binning process using the binning module of metaWRAP v1.3.2 [64] (parameters: --metabat2 --maxbin2 --concoct). Subsequently, a set of bins was used for Bin_refinement module (parameters: -c 50 -x 10) within metaWRAP. After bin refinement, dRep v3.4.0 [65] (parameters: -comp 50, -con 10, -sa 0.95) was used for dereplication at 95% average nucleotide identity (ANI), resulting in a non-redundant set of 1,006 species-level metagenome-assembled genomes (MAGs). Taxonomy of MAGs was performed by GTDB-Tk v2.3.2 [66] based on the Genome Taxonomy Database (GTDB, http://gtdb.ecogenomic.org) using R214 reference package. Using GTDB-Tk, 120 bacterial and 53 archaeal marker genes were identified and processed, which included alignment, concatenation, and trimming. The host genomes were utilized for the construction of domain-specific phylogenetic trees through IQ-TREE v2.2.0.3 [67, 68], utilizing best-fit models and conducting 1000 bootstrap iterations. The phylogenetic trees of bacteria and archaea were visualized using Interactive Tree Of Life (iTOL,v6) [69].

Detection of antiviral systems in MAGs was carried out using DefenseFinder v1.0.9 based on MacSyFinder models according to MacSyfinder rules (parameters: -dbtype gembase) [18, 70]. The metabolic functions of MAGs were annotated using METABOLIC v4.0 [71] (parameters: default) and DRAM v1.3.5 [72] (parameters: default).

### Identification and classification of vOTUs

Potential viral contigs were identified from metagenome assemblies (contigs > 10 kb) using geNomad v1.7.0 [73] (parameters: end-to-end), DeepVirFinder v1.0 [74] (parameters: default), Virsorter2 v2.2.3 [75] (parameters: default), and VIBRANT v1.2.1 [76] (parameters: default). The union of potential viral contigs was used for subsequent analysis. To assess the completeness and contamination of the identified contigs from the four methods mentioned above, we employed CheckV v1.0.1 [77] (database version v1.5). Genomes with an estimated completeness of 50% or greater were then grouped into species-level viral operational taxonomic units (vOTUs) using the parameters 95% average nucleotide identity and a minimum aligned fraction of 85% (https://bitbucket.org/berkeleylab/checkv/src/master/) provided in CheckV. Finally, a total of 1,376 species-level vOTUs were identified. The open reading frames (ORFs) of each vOTU were predicted by Prodigal v2.6.3 [78] (parameters: -p meta).

For taxonomic assignments of viruses, four approaches were adopted: (i) The geNomad v1.7.0 (parameters: end-to-end) was employed for the taxonomic assignment of viral genomes. (ii) Viral genomes were classified using a taxonomy based on the International Committee on Taxonomy of Viruses (ICTV) of IMG/VR (v4) database [79]. The methodologies were guided by the scripts provided in the ViWrap toolkit [80]. Specifically, ORFs of each vOTU were searched against the IMG/VR (v4) database using Diamond v2.0.8.146 [81] (parameters: --evalue 0.00001 --query-cover 50 -- subject-cover 50). The taxonomy for each ORF was determined based on the top hit from the alignment results. Each vOTU was assigned to the lowest taxonomic rank of >50% of the annotated proteins. (iii) vConTACT2 v0.11.3 was utilized to annotate viral genomes based on the NCBI Reference Database. According to the International Committee on Taxonomy of Viruses (ICTV) classification, viruses within the same clusters are approximately equivalent to those within the same genus [82]. For vConTACT3 v3.0.0b26, based on the v211 reference database, classification was specifically conducted at the genus level for vOTUs that were fully clustered (https://bitbucket.org/MAVERICLab/vcontact3/src/master/). (iv) 77 single-copy genes were used for constructing phylogenetic trees of *Caudoviricetes* [83]. The vOTUs in this study were classified to the family level if they were assigned to a known clade within NCBI RefSeq.

To compare the 1376 vOTUs with those from other environments, viral contigs from public databases were collected: (i) acid mine drainage [84] (n = 15032); (ii) cold seep [85] (n = 2885); (iii) waste water [49] (n = 2885); (IV) soil [86] (n = 1907); (V) seawater [87] (n = 195728). The protocol in protocols.io (https://www.protocols.io/view/applying-vcontact-to-viral-sequences-and-visualizi-kqdg3pnql25z/v5) were selected to analyse vOTUs from different environments. Specifically, ORFs of each vOTU were predicted by Prodigal and the predicted protein sequences were clustered using vConTACT2 v0.11.3 (parameters: --db ’ProkaryoticViralRefSeq94-Merged’ --pcs-mode MCL --vcs-mode ClusterONE --rel- mode Diamond). Afterwards the genome gene-sharing profiles were visualized by the gene-sharing network in Cytoscape v3.10.1 [88].

To enhance understanding of *Caudoviricetes* of this study, *Caudoviricetes* viruses were downloaded from the NCBI Reference Database to the family level. 77 defined *Caudoviricetes* markers against the vOTUs’ protein-coding sequences were searched using a profile Hidden Markov Model (HMM) as described [83]. Subsequently, the highest-ranking HMM matches were aligned with the profile HMMs corresponding to the 77 markers, with each marker gene being aligned and trimmed independently. Alignment was conducted using MUSCLE v3.8.1551 [89] (parameters: default), and trimming was performed with trimAl v1.4.rev15 [90] (parameters: default); the remaining regions had gaps less than 50%. Genomic sequences were concatenated, and gaps were filled using a Python script. Genomes representing more than 5% of the amino acid content in the total alignment length were selected for further analysis. A phylogenetic tree of concatenated proteins was constructed from the aligned sequences using FastTreeMP v2.1.10 under the automatic model option available online (https://www.phylo.org/portal2/) [91]. Phylogenetic trees were visualized with iTOL v6.

### Viral lifestyle prediction

For lifestyle prediction, three methods were employed. (i) BACPHLIP v0.9.6 [93] (minimum score of ≥0.8) and VIBRANT v1.2.1 (parameters: default) were used to infer the lifestyle of complete viral genomes. (ii) CheckV v1.0.1 was used to predict the lysogenic lifestyle of viral genomes by detecting provirus boundaries. (iii) ORFs were functionally annotated by eggNOG-mapper v2.1.9 [94] (parameters: m diamond). Sequences containing provirus integration sites or integrase genes were considered to be lysogenic.

### Abundances of MAGs and vOTUs

Reads per kilobase per million mapped reads (RPKM) values were used to represent the relative abundance of MAGs and vOTUs. CoverM v0.6.1 [95] (parameters: coverm contig --trim-min 0.1 --trim-max 0.90 --min-read-percent-identity 0.95 --min-read-aligned-percent 0.75 -m rpkm) was used to calculate the RPKM values for the MAGs and vOTUs.

### Virus-host linkages of viruses

A total of 1,006 bacterial and archaeal MAGs identified in this study and iPHoP_db_Sept21_rw were used to construct a host reference database. Virus‒host linkages were predicted based on multiple strategies using the iPHop v1.3.2 [96] pipeline: (i) blastn to host genomes: Blastn v2.12.0+ [97] was utilized for comparing viral genomes with databases. All hits with ≥ 80% identity and ≥ 500bp were considered. Hits covering ≥ 50% of the "host" contig length are ignored. (ii) Blastn to CRISPR spacer: CRISPR spacer databases of iPHoP_db_Sept21_rw were predicted in advance. Blastn v2.12.0+ was utilized for comparing viral genomes against CRISPR spacer databases. The selection criteria were focused on hits that aligned with spacers of at least 25 nucleotides in length, with fewer than 8 total mismatches, and a custom complexity score below 0.6. Additionally, all hits with up to 4 mismatches were considered in the analysis. (iii) RaFAH [98]: RaFAH v0.3 was used to predict viral genomes with default settings. The Predicted_Host_Score was employed as the primary metric for these predictions. (iv) WisH [99]: Viral genomes were compared to the WIsH iPHoP_db_Sept21 database with WIsH v1.0 and a maximum p-value of 0.2; (v) VHM: Using the VirHostMatcher-Net (version of July 2021), the input viral genomes were compared against the VirHostMatcher iPHoP_db_Sept21 database. For each hit identified, the s2* similarity score was used as the metric for evaluation. (vi) PHP: Viral genomes were compared with those of the PHP iPHoP_db_Sept21 database using the PHP tool (version of July 2021). The PHP score served as the evaluative metric for each identified hit.

To identify potential archaeal viruses, prior text-based tool MArVD2 [92] (parameters: default) was employed for discerning archaeal viruses within the vOTUs. Only vOTUs with Archaeal Virus Probability of 0.8 or higher were retained.

### Identification of auxiliary metabolic genes, anti-defense genes, antibiotic resistance genes and metal resistance genes

For identification of AMGs, Virsorter2 v2.2.3 (parameters: --prep-for-dramv) were used to produce files for DRAM v1.3.5 (“virome” model). AMGs with auxiliary scores of 1-3 and flags of M were selected for further manual curation. In manual curation, illegal AMGs (such as nucleotide metabolism, ribosomal proteins, glycosyl transferases, ‘Organic Nitrogen’ category) were removed. Afterwards, a detailed examination of the upstream and downstream regions of the AMGs was conducted. Three-dimensional structures of AMGs were predicted by ColabFold v1.5.5 using AlphaFold2 [100–104]. Phylogenetic tree of AMG *napA* was constructed using IQ-TREE v2.2.0.3 with the best-fit models and 1000 bootstrap iterations after aligning sequences by MUSCLE v3.8.1551 (parameters: default). Phylogenetic tree was visualized with the Interactive Tree Of Life (iTOL, v 6). The comparison of AMG *napA* protein structures was based on two methods: (i) PyMOL v2.5.0a0 was used to calculate the RMSD values between proteins [105]. (ii) The TM-score, available through an online tool (https://zhanggroup.org/TM-score/), was used to assess similarities in protein structure topology. The HDOCK web platform was used to dock *napA* and *napB* proteins from viruses, yielding an empirical confidence rating of 0.7, indicating binding between the two proteins [106]. The highest-ranked model was selected to show interactions between proteins [107].

For the identification of anti-defense genes, dbAPIS [108] was selected as a reference database. Diamond v2.0.8.146 (parameters: -e 1e^-10^ --id 30 --max-target-seqs 1) was used to annotate anti-defense genes from viral sequences following the protocol provided by dbAPIS (https://bcb.unl.edu/dbAPIS/downloads/readme.txt).

For the identification of ARGs and MRGs, two methods were employed: (i) Viral sequences were searched against Comprehensive Antibiotic Resistance Database [109] (CARD, v3.2.8) and Antibacterial Biocide & Metal Resistance Genes Database [110] (BacMet v2.0) using BLAST 2.5.0+ with e-value ≤ 10^−5^ and identity ≥ 50%. (ii) The geNomad v1.7.0 (parameters: end-to-end) was employed to annotate viral ARGs and MRGs. To compare homology between viral MRGs, ARGs and host MAGs, BLAST 2.5.0+ (parameters: blastn -perc_identity 70 -qcov_hsp_perc 80) was used.

To understand the function of viruses with lytic cycles, the HHpred online server was used to annotate the structural proteins of viruses with lytic cycles infected with corrosive microbes via structure-based HHpred searches [111, 112].

### Statistical analyses and plotting

Statistical analyses were performed using R v4.3.1. The alpha and beta diversity of the communities were calculated using the vegan package v2.6–4. The alpha diversity indices included the Shannon, Chao1, and observed species (Obs) indices. Wilcoxon tests were used to compare alpha diversity among different materials.

To calculate beta diversity, Bray‒Curtis dissimilarities were derived from the OTU tables. Both principal coordinate analysis and nonmetric multidimensional scaling were utilized for dimensionality reduction and analysis. Correlations between the relative abundance of viruses and their hosts were assessed using Spearman’s rank correlation coefficient, where the strength and direction of the monotonic relationship were characterized by the correlation coefficient and p value.

Gene clusters and bar charts were drawn using ChiPlot (https://www.chiplot.online/). The workflow was drawn using Drawio v23.1.5 online (https://app.diagrams.net/). A Sankey plot was constructed via the Pavian [113] (https://fbreitwieser.shinyapps.io/pavian/).

## Authors’ contributions

R.Z. and X.D. designed the research; R.Z., X.D. and Y.Z. gave important suggestion about data analysis; C.L. performed data analysis, integration and wrote the manuscript and prepared figures and tables. R.Z., X.D. S.J., Y.Z., W.S., Y.P. and Y.H.reviewed drafts of the manuscript. All authors approved the final manuscript.

## Availability of data and materials

Data of metagenome used in this study is provided in Table S1.

## Acknowledgements

This study relied on the publicly available genomic data by previously published research. We extend our sincere gratitude to all the participants, organizations, and funding agencies responsible for the collection and analysis of the data used in our study.

## Ethics approval and consent to participate

Not applicable.

## Consent for publication

Not applicable.

