## Supplemental figures for "Viral diversiy within marine biofilms and their potential roles in microbially influenced corrosion"

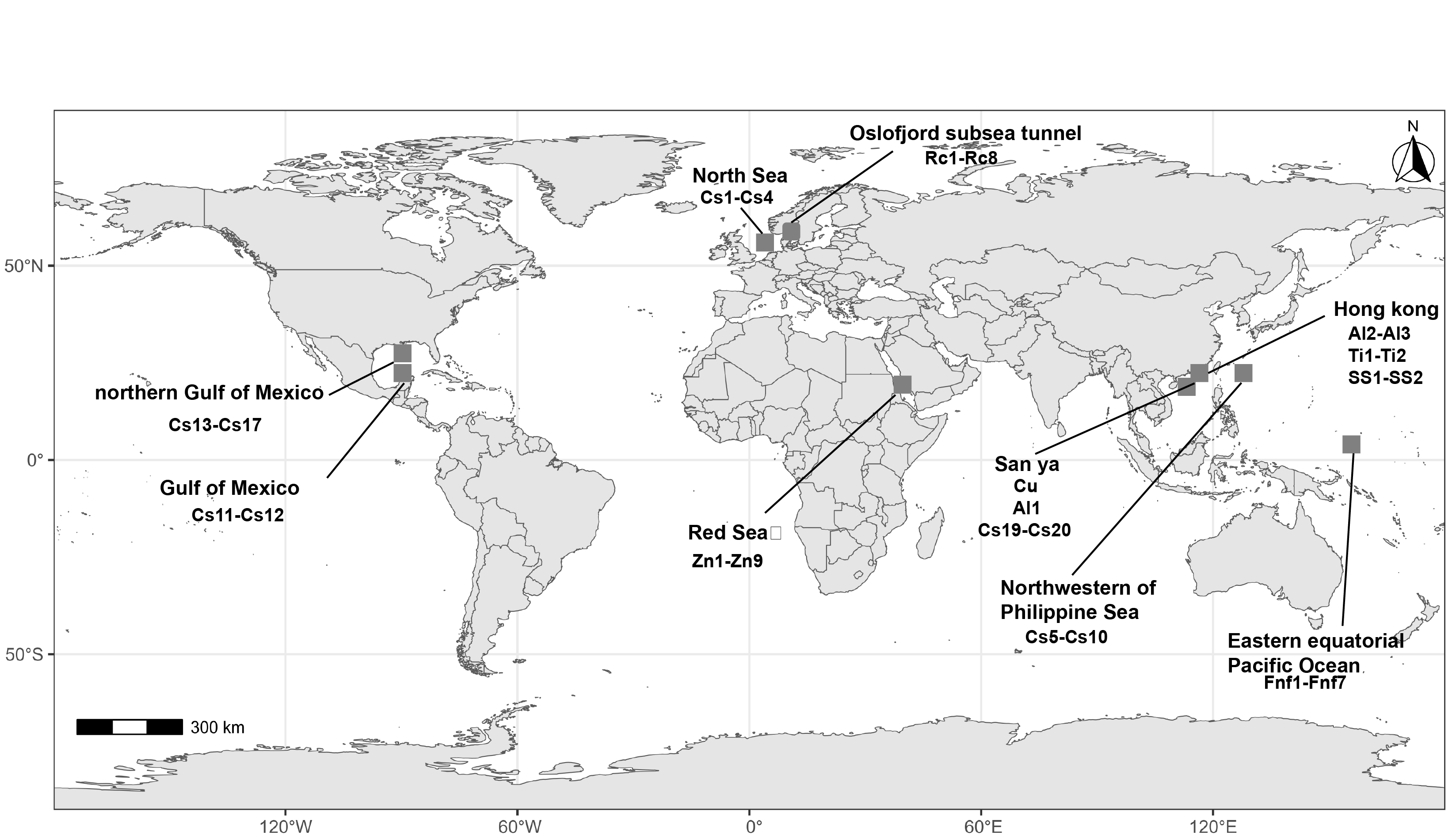

**Figure S1. Geographical distribution of collected metagenomic data.** The location where the material was collected includes the abbreviation of the sample name. A sample cultured in the laboratory is not shown in this figure.

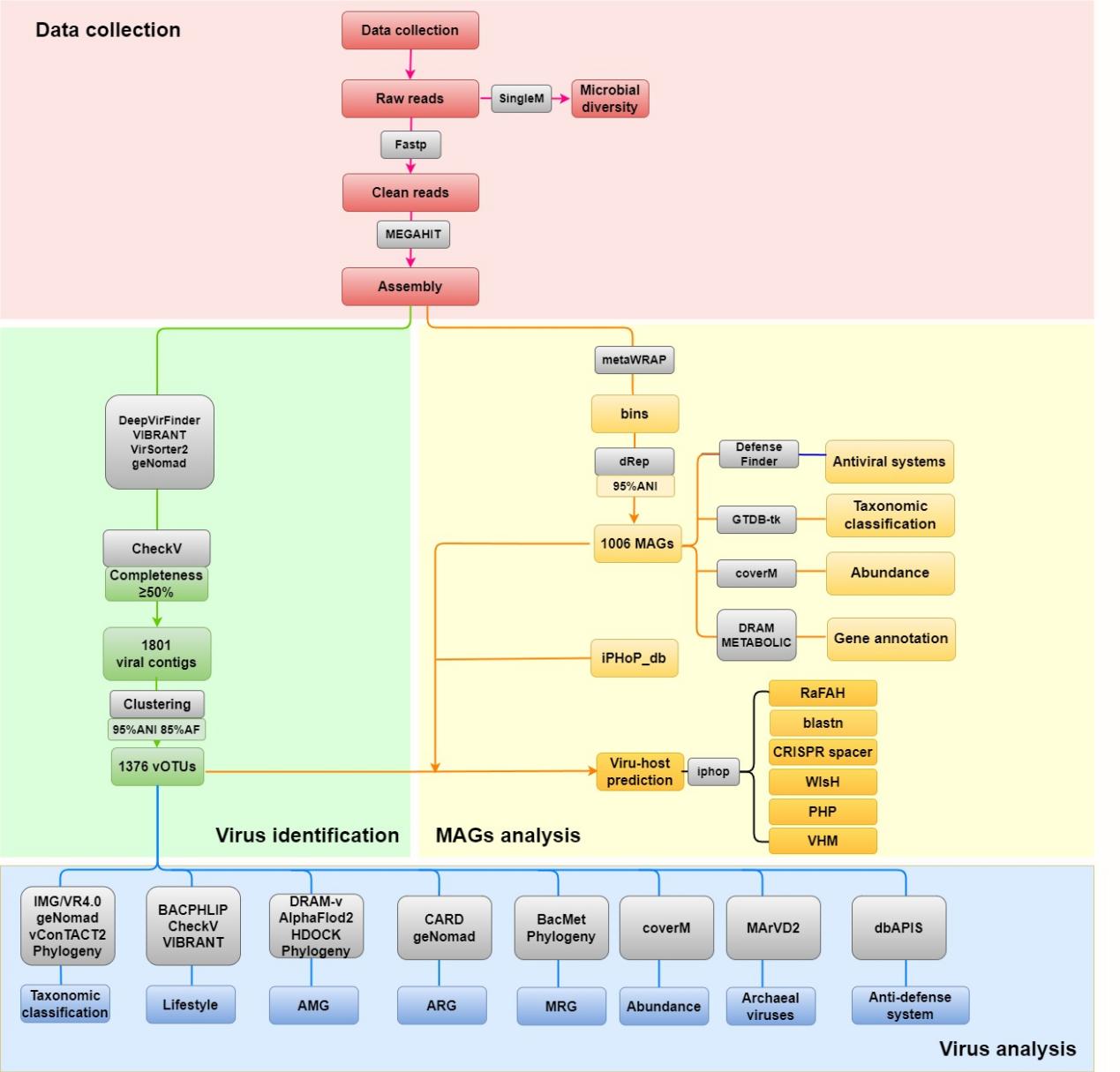

**Figure S2.** **Workflow for viral analysis in this study.** Red: Data Collection; Green: Virus Identification; Yellow: MAGs Analysis and Host prediction Blue: Virus analysis

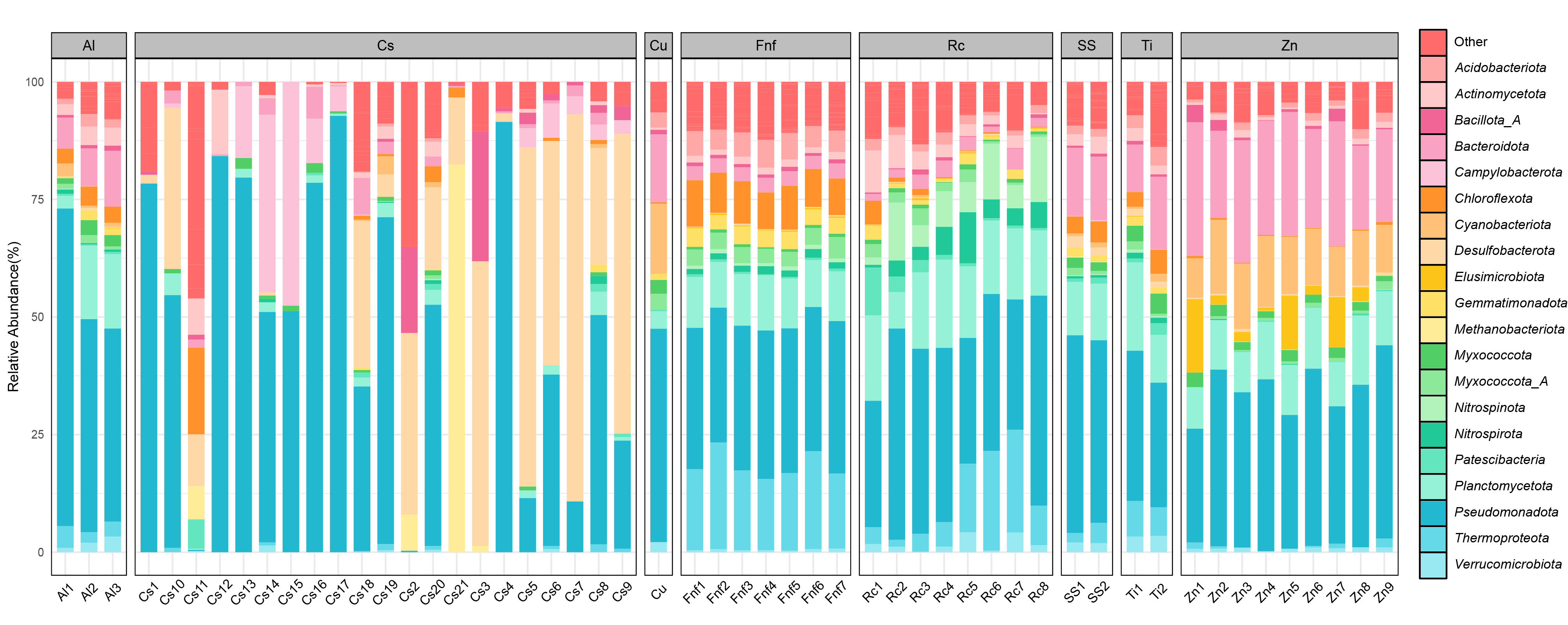

**Figure S3. The relative abundance of prokaryotic taxa at the phylum level across different materials.** Comparison of taxonomy in prokaryotic community among 53 samples from different types of materials based on *rplB*. Bar chat shows the top 20 phyla by relative abundance.

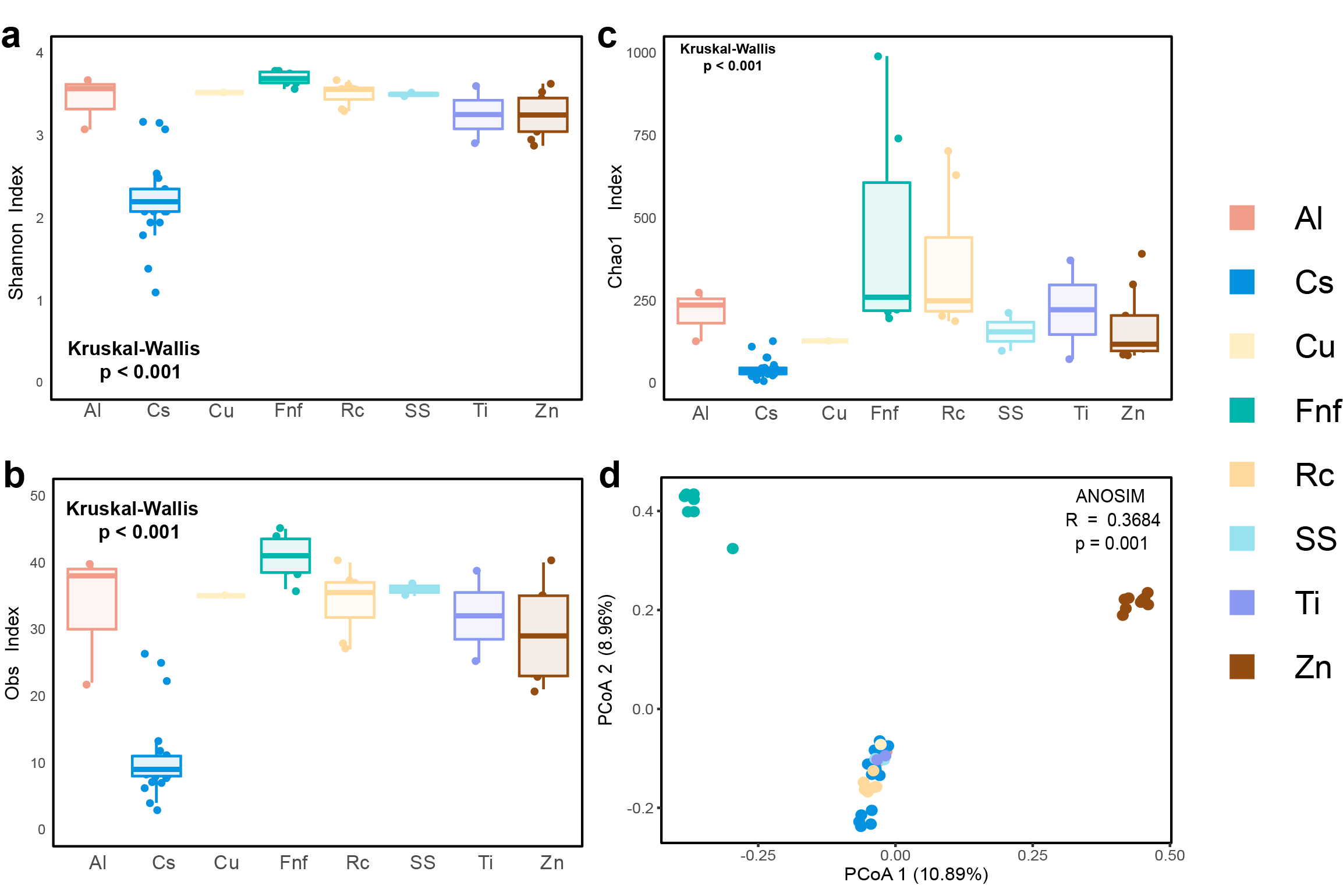

**Figure S4. Comparison of prokaryotic community diversity between different materials. a** Shannon indices of prokaryotic community diversity from different materials. **b** Obs indices of prokaryotic community diversity from different materials. **c** Chao1 indices of prokaryotic community diversity from different materials. **d** PCoA analysis of a Bray-Curtis dissimilarity matrix from OTUs. ANOSIM was applied to test the difference in communities between different materials.

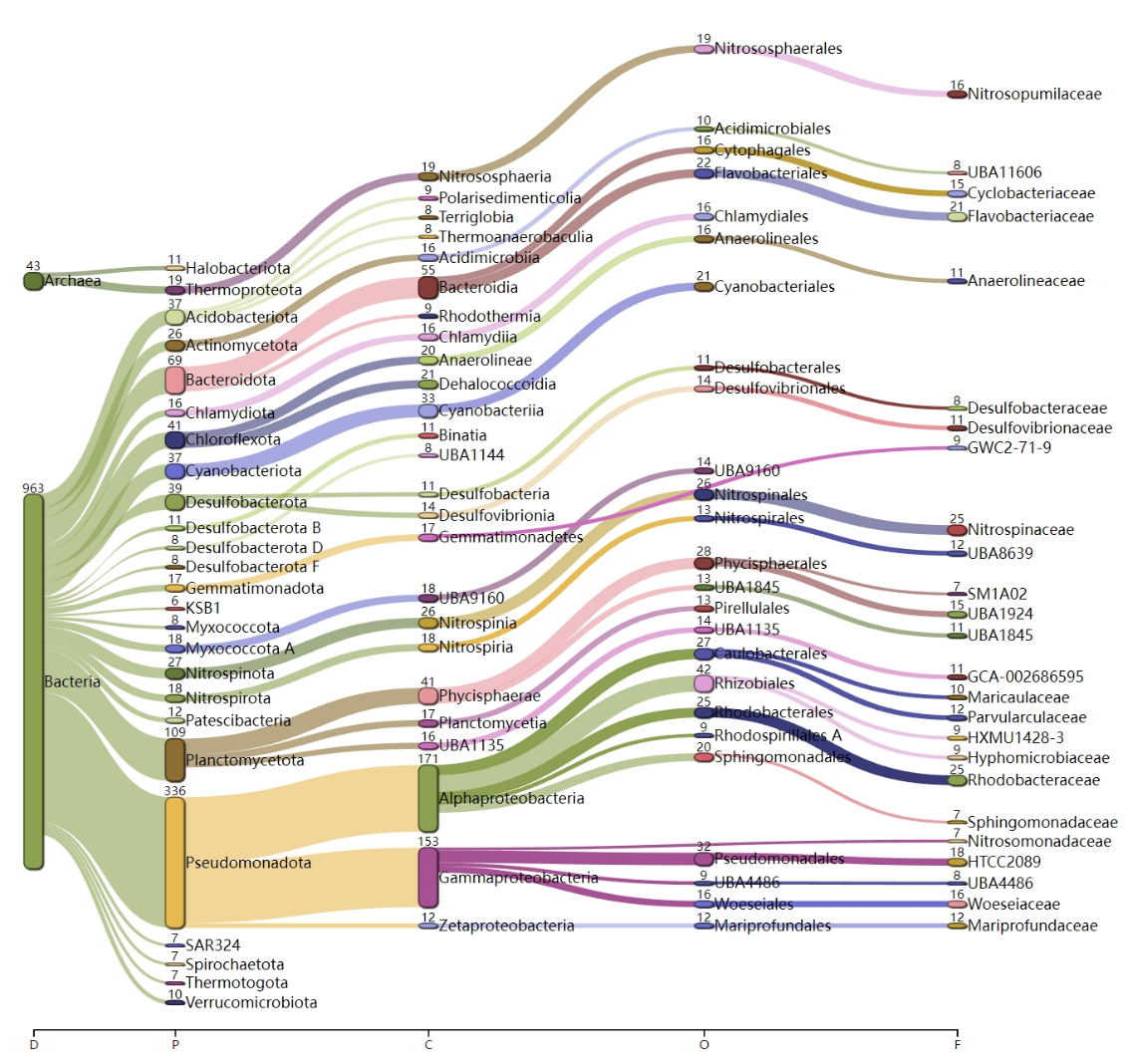

**Figure S5. Classification of MAGs recovered from different materials.** Sankey diagram illustrating the taxonomic assignment based on GTDB taxonomy, displaying archaeal and bacterial MAGs across various taxonomic levels. The diagram shows the top 25 taxa with the highest number of MAGs at each level.

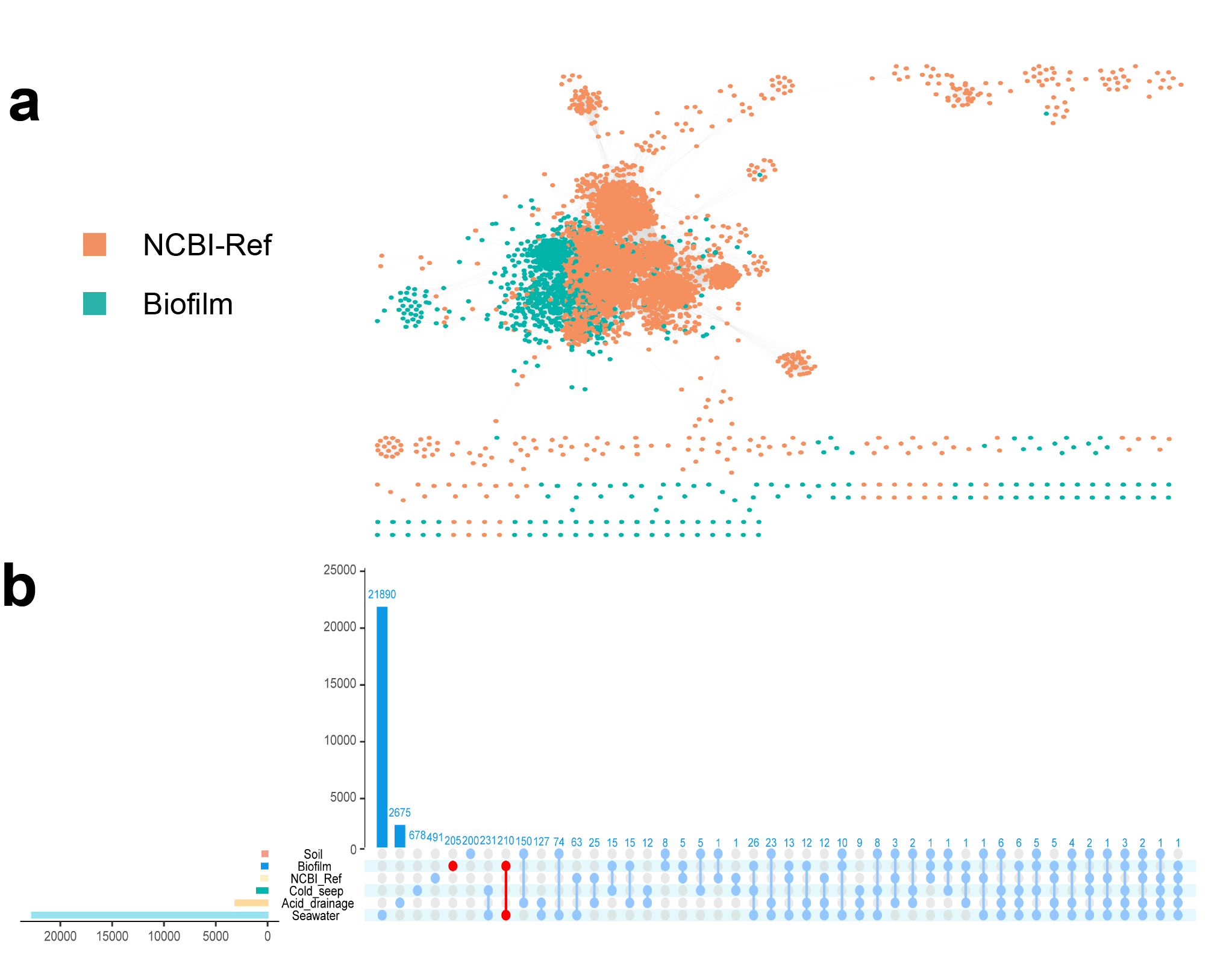

**Figure S6. Classification of viruses from biofilms and comparison with viruses from other environment. a** Gene-sharing network of viral sequence from biofilm and RefSeq viral genomes. Each node represents a viral genome and each edge represents a shared protein cluster. **b** Comparative analysis of viruses from biofilm with RefSeq and other viruses in other environments. Upset diagram of shared viral clusters among 5 datasets and RefSeq. The two most abundant viral protein clusters from biofilm are highlighted in red.

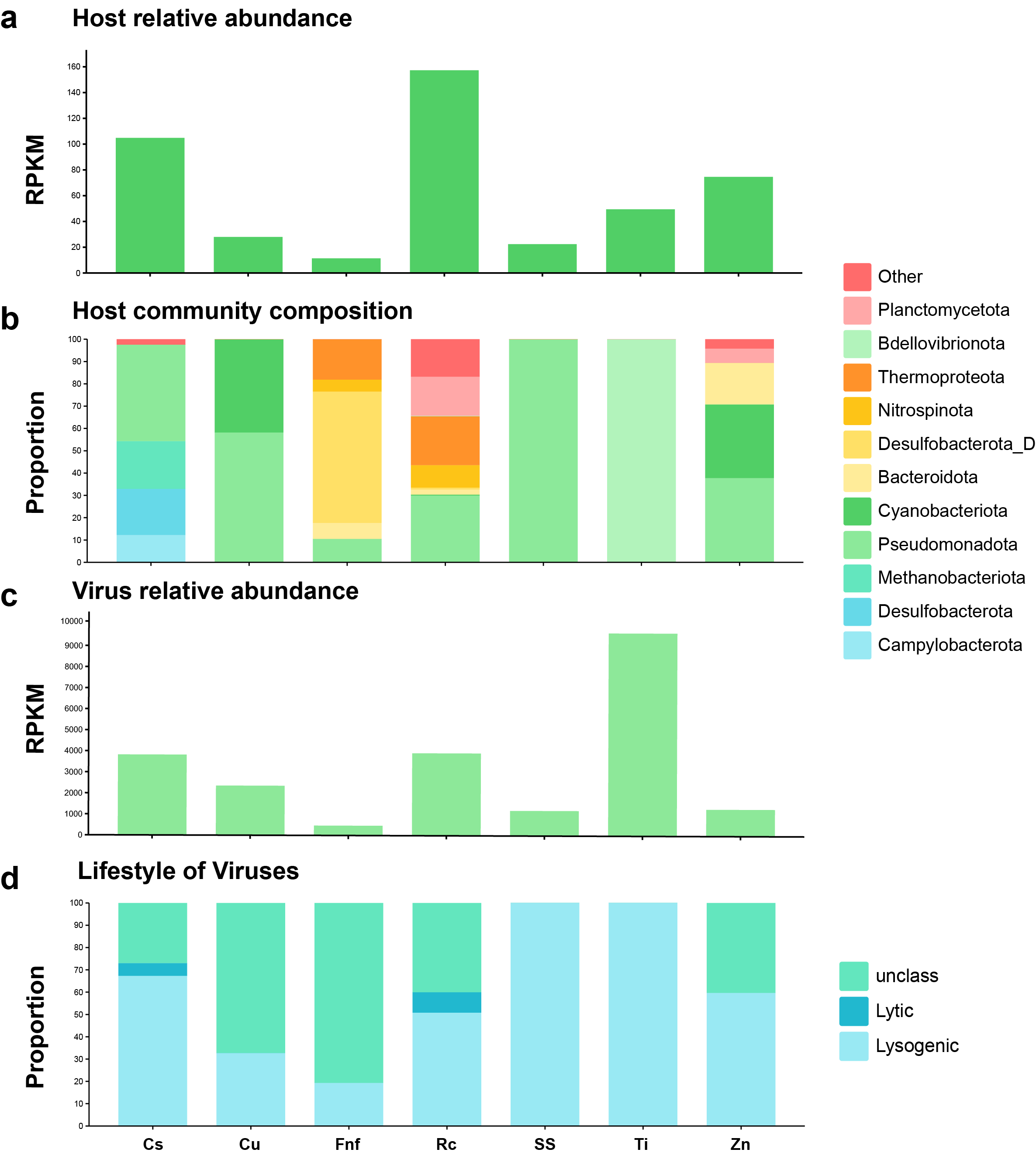
**Figure S7. The relative abundances of viruses and hosts across different materials.** **a** The relative abundances of host based on RPKM values. **b** The proportion of host taxa at the phylum level. **c** The relative abundances of viruses based on RPKM values. **d** The proportion of virus lifestyle.

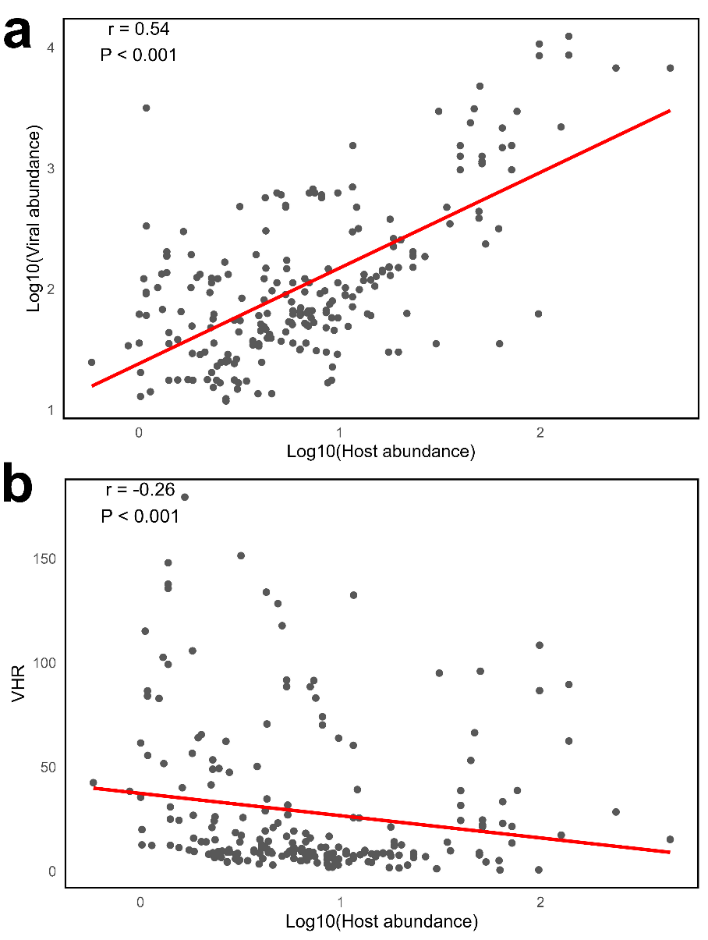

**Figure S8. The linkages of relative abundance between viruses and hosts.** a Correlation between the relative abundance of viruses with lysogenic cycles and hosts. The regression line is represented in red. b Correlation between the relative abundance of viruses and virus-host abundance ratios (VHR). The regression line is represented in red.

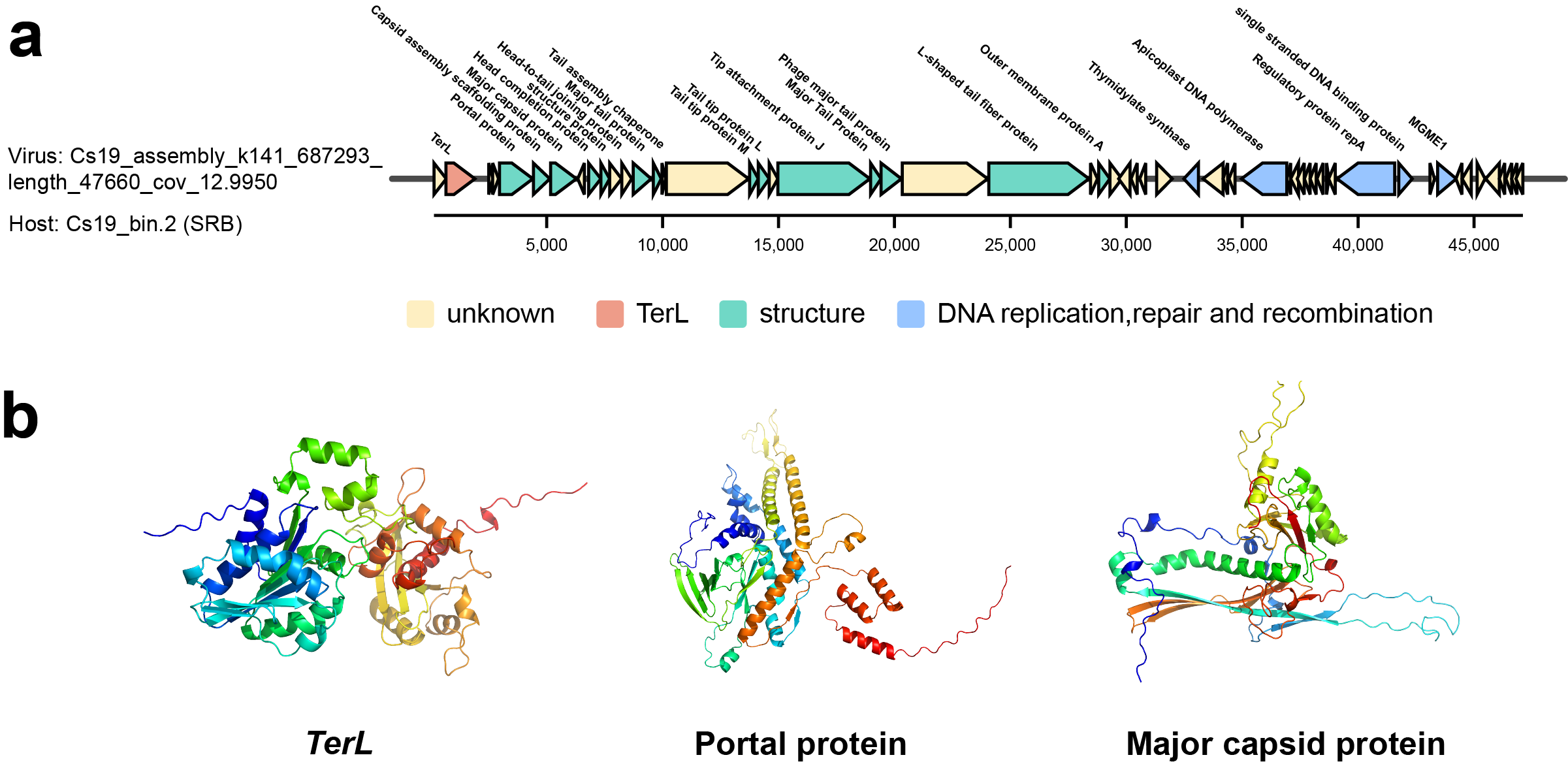

**Figure S9. The genomic context and protein structure of virus with a lytic cycle infected on sulfate-reducing prokaryotes.** *TerL* are marked in red, structure genes are marked in cyan, DNA replication, repair, and recombination genes are marked in blue, genes with unknown function are marked in yellow. Protein structure colors are automatically assigned using PyMOL.

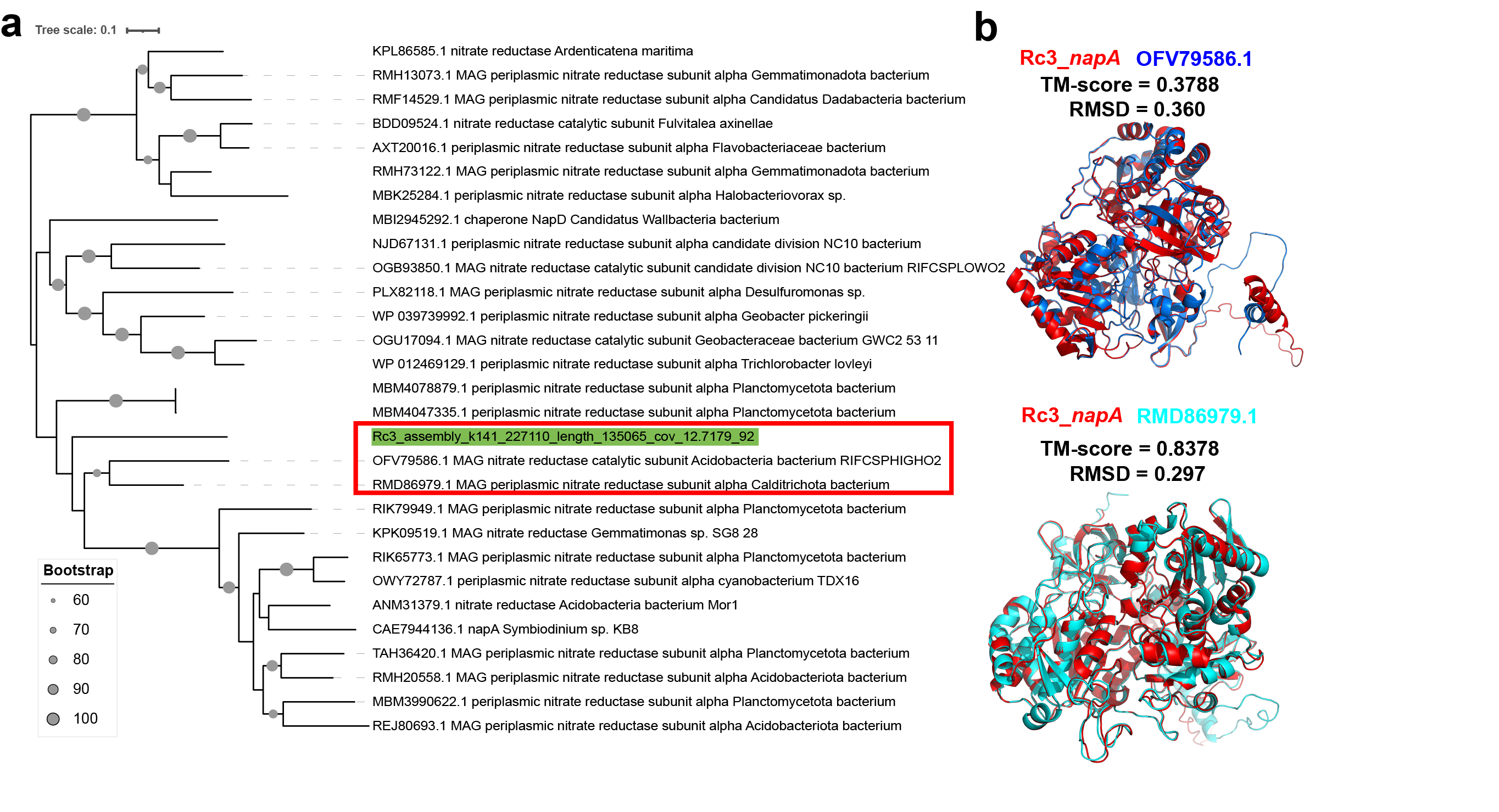

**Figure S10. Relationship between *napA* from virus in this study and RefSeq bacterial proteins.** **a** Maximum-likelihood phylogeny of *napA*. Viral AMGs and the two nearest sequences are highlighted in red boxes. **b** Pariwise comparison of *napA* between viral AMG and bacterial protein from RefSeq. The viral AMGs are in red, while the bacterial proteins are in blue and cyan. TM-score was calculated on the Zhang Lab (https://zhanggroup.org/TM-score/), and RMSD was calculated using PyMOL.

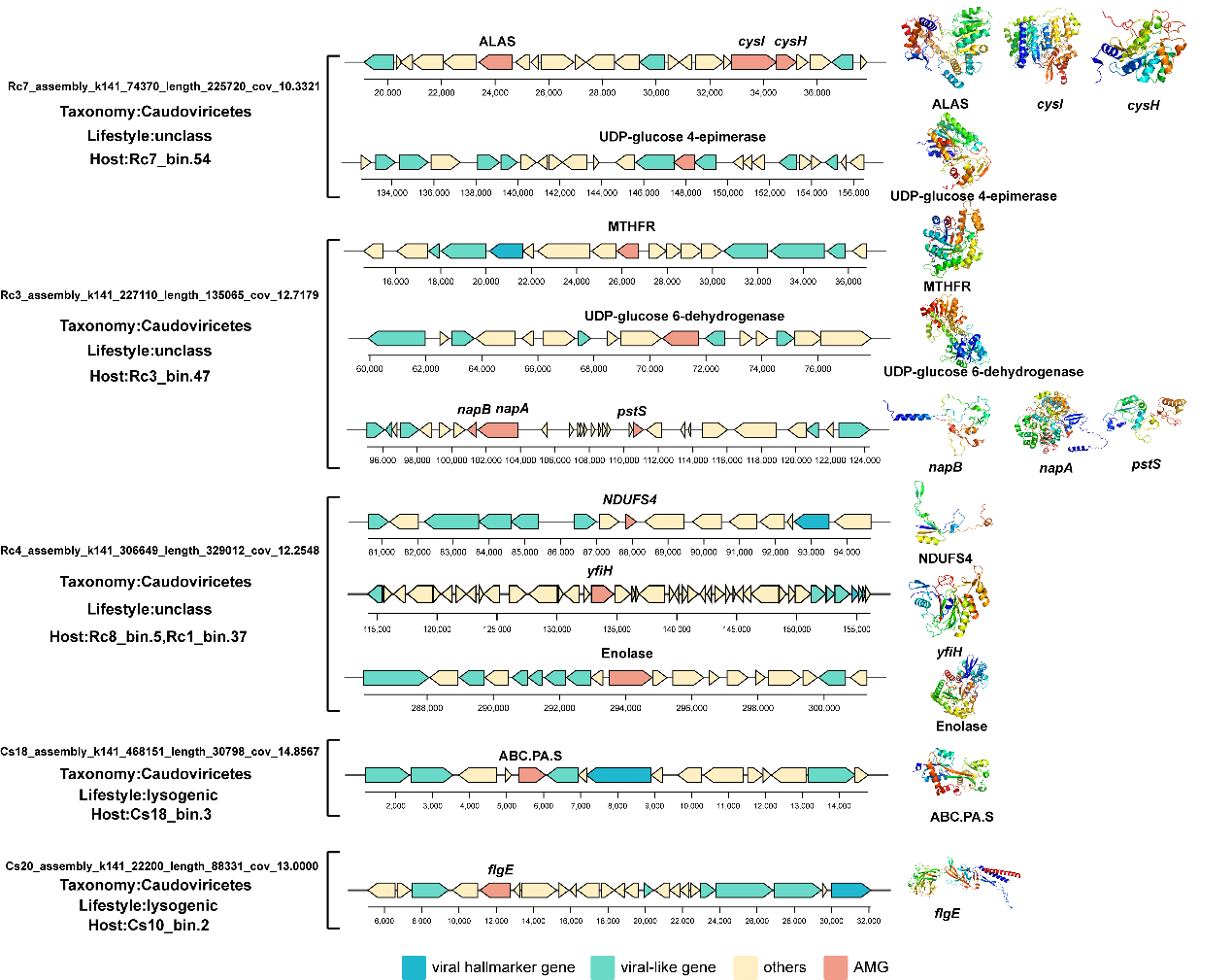

**Figure S11.** **The genomic context and protein structure of selected viral auxiliary metabolic genes.** AMGs are marked in red, viral-like genes are marked in cyan, viral hallmark genes are marked in blue, and other genes are marked in yellow. Detailed information on AMGs can be found in Table 14. Protein structure colors are automatically assigned using PyMOL.

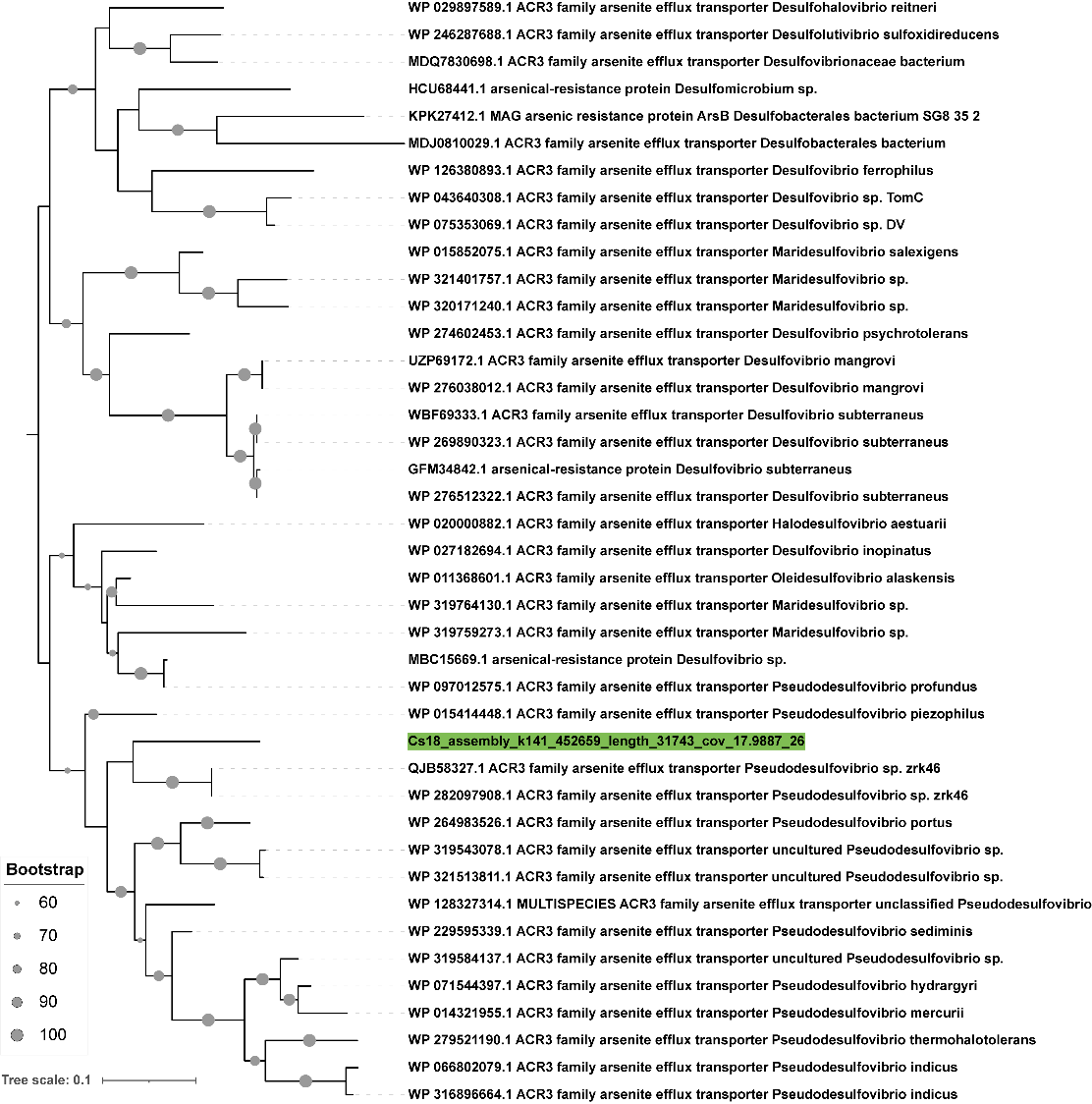

**Figure S12. Relationship between *arsB* from virus in this study and RefSeq bacterial proteins.** Maximum-likelihood phylogeny of *arsB*. Viral AMGs are highlighted in green.

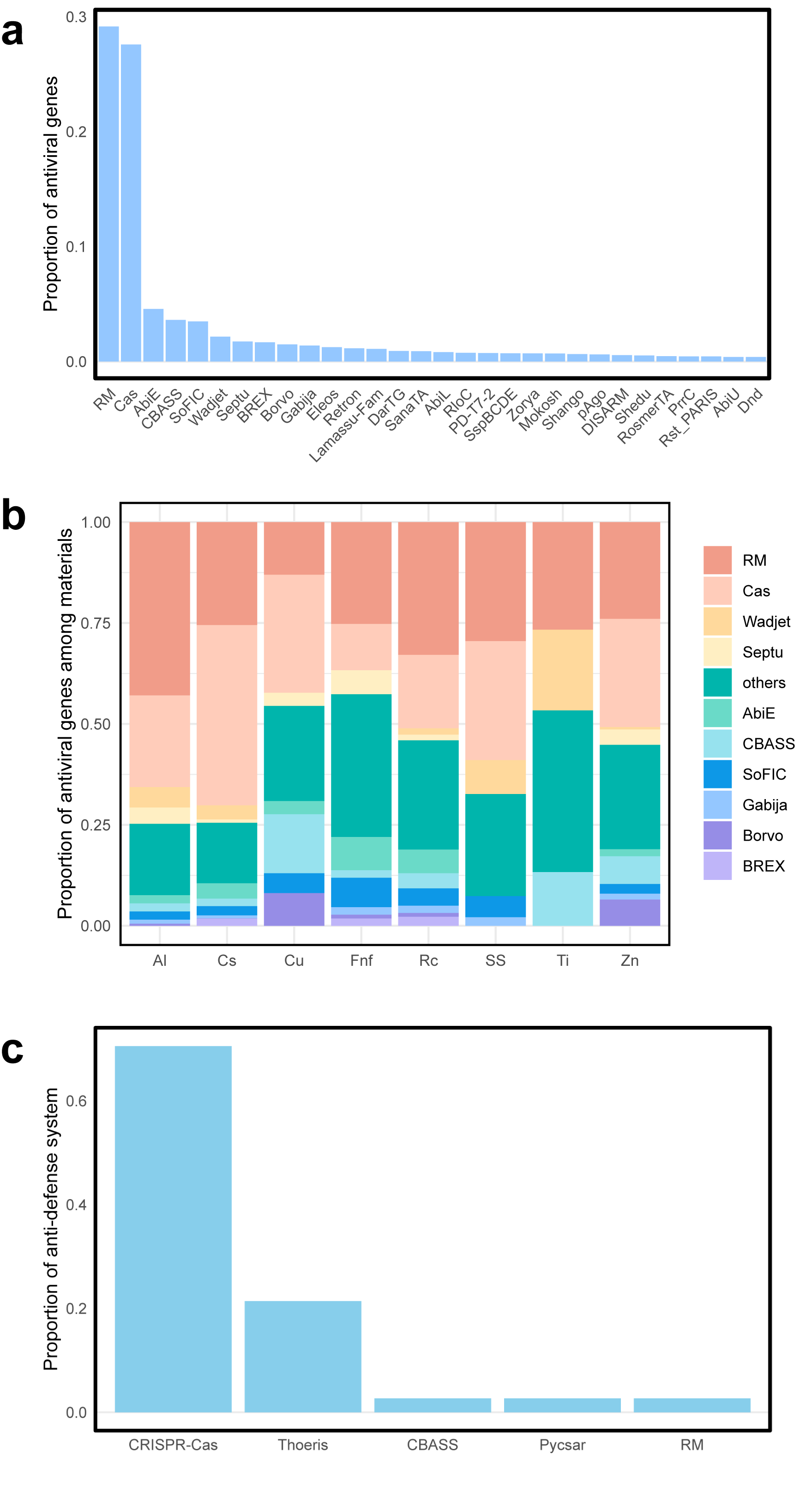

**Figure S13. Anti-viral systems and anti-defense systems found in prokaryotic genomes and viral genomes.** **a** The proportion of anti-viral genes in each anti-viral system. **b** The distribution of anti-viral genes across different materials. **c** The proportion of anti-defense genes in each anti-defense system.
